# iMUT-seq: high-resolution mapping of DSB mutational landscapes reveals new insights into the mutagenic mechanisms of DSB repair

**DOI:** 10.1101/2021.12.08.471781

**Authors:** Aldo S. Bader, Martin Bushell

## Abstract

DNA double-strand breaks (DSBs) are the most mutagenic form of DNA damage, and play a significant role in cancer biology, neurodegeneration and aging. However, studying DSB-induced mutagenesis is currently limited by the tools available for mapping these mutations. Here, we describe iMUT-seq, a technique that profiles DSB-induced mutations at high-sensitivity and single-nucleotide resolution around endogenous DSBs spread across the genome. By depleting 20 different DSB-repair factors we defined their mutational signatures in detail, revealing remarkable insights into the mechanisms of DSB-induced mutagenesis. We find that homologous-recombination (HR) is mutagenic in nature, displaying high levels of base substitutions and mononucleotide deletions due to DNA-polymerase errors, but simultaneously reduced translocation events, suggesting the primary role of HR is the specific suppression of genomic rearrangements. The results presented here offer new fundamental insights into DSB-induced mutagenesis and have significant implications for our understanding of cancer biology and the development of DDR-targeting chemotherapeutics.

## INTRODUCTION

DNA double-strand breaks (DSBs) are highly mutagenic lesions that commonly result in genomic base substitutions, deletions and chromosomal rearrangements. DSBs occur in our genomes everyday due to exogenous sources, such as radiation, as well as endogenous sources, such as transcription-replication conflicts. As a result, effective DSB-repair (DSBR) is critical for genome maintenance to prevent the toxic accumulation of DSB-induced mutations, which are commonly associated with diseases, most notably cancer. Defects in DSBR are commonly associated with cancer-prone syndromes as they promote increased mutagenesis, driving carcinogenesis as well as cancer progression.

There are two major DSBR pathways; non-homologous end-joining (NHEJ) and homologous recombination (HR) operate via distinct mechanisms to provide comprehensive repair. NHEJ is initiated by binding of the KU70-80 heterodimer to the exposed DNA at the DSB together with the DNA-PK catalytic subunit, DNA-PKcs, to form DNA-PK (Blackford and Jackson, 2017) which phosphorylates and recruits downstream targets (Anisenko *et al*, 2020; Mohiuddin and Kang, 2019). End processing factors such as Artemis and PNKP are then recruited to ensure the broken DNA ends are ready for ligation (Anisenko *et al*, 2020; Weterings and Chen, 2008). Recruitment of XRCC4 and XLF acts as a scaffold around the DNA which facilitates further recruitment of NHEJ factors (Ropars *et al*, 2011; Roy *et al*, 2012; Mahaney *et al*, 2013; Andres *et al*, 2012), such as polymerase-λ (POLL), and Ligase IV (LIG4) that carries out the final ligation to complete repair (Fan and Wu, 2004; Pryor *et al*, 2015). This pathway for DSBR provides a rapid response that is broadly applicable to most DSBs; however, NHEJ lacks any checks on repair fidelity, and therefore is known to result in deletions at the break and also chromosomal rearrangements (Richardson and Jasin, 2000; Mao *et al*, 2008).

During HR, initially the MRN complex (MRE11-RAD50-NBS1) is required for short-range DNA end resection that occurs bi-directionally at DSBs (Garcia *et al*, 2011; Shibata *et al*, 2014; Reginato, Cannavo and Cejka, 2017) and the ATM kinase also phosphorylates a number of downstream factors to both activate and recruit them to breaks (Blackford and Jackson, 2017). BRCA1 maintains and promotes resection, while long-range resection enzymes such as EXO1 and the BLM-DNA2 complex then extend resection kilobases from the break (Grabarz *et al*, 2013; Gravel *et al*, 2008; Karanja *et al*, 2012; Nimonkar *et al*, 2011; Sturzenegger *et al*, 2014). RAD51 forms a filament around this DNA overhang in a BRCA2-dependent manner (Moynahan, Pierce and Jasin, 2001; Han *et al*, 2017) facilitating invasion of the intact sister-chromatid, allowing for re-polymerisation of the DNA over the break. DNA polymerisation during HR is known to occur via multiple polymerases including polymerase-δ (Li, X. *et al*, 2009; McVey *et al*, 2016) and polymerase-ε (McVey *et al*, 2016). HR therefore provides a method of DSBR that maintains DNA fidelity by using the sister-chromatid as a template.

Although HR is theoretically higher fidelity, recent reports have suggested mutations still occur during HR due to polymerase error and repetitive sequences causing misalignment of the resected DNA to the sister-chromatid (Yang *et al*, 2008; Hicks, Kim and Haber, 2010; Guirouilh-Barbat *et al*, 2014). In addition, the requirement of a sister-chromatid limits HR to the S- and G2-phases of the cell-cycle, causing most DSBR to still be carried out by NHEJ.

There are currently a number of methods for the investigation of DSB-induced mutations, which can be divided into whole genome sequencing (WGS) or exogenous reporter-based approaches. Whereas WGS provides a direct measure of genomic mutations under different conditions, since any damage induced, e.g. via irradiation or chemical induction such as etoposide, will be randomly acquired across the genome, you cannot gain an insight into how mutations are introduced relative to the site of damage. In addition, the size of the human genome causes high read-depth to be impractical with modern sequencing techniques, and therefore very high levels of DNA damage need to be used in order to observe the mutagenic signatures (Kucab *et al*, 2019). Reporter systems for DSB-induced mutations use either restriction enzymes or CRISPR-Cas9 to induce DSBs at defined reporter sequences that have been added to the genome (Ahrabi *et al*, 2016; Hussmann *et al*, 2021; Schep *et al*, 2021). These approaches can therefore map mutations around the DSB at nucleotide resolution, but not at regions endogenous to the genome which may limit their applicability. It is also common to use high levels of DNA damage in these systems to enrich for mutations, often expressing the restriction or CRISPR enzymes for multiple days (Ahrabi *et al*, 2016; Hussmann *et al*, 2021; Schep *et al*, 2021), which will cause high levels of damage over multiple rounds of repair and will significantly alter mutation signatures. There is therefore a need for a technique that provides the quantification of mutations around known, endogenous DSB loci and that can provide very high sensitivity to detect rare mutation events.

Here, we describe our newly developed iMUT-seq technique that uses an inducible DSB system to introduce DSBs across the genome at defined endogenous loci followed by targeted next-generation sequencing (NGS) to profile all DSB-induced mutations at single-nucleotide resolution with extremely high sensitivity around these DSBs up to 100bp from the break. We then systematically depleted or inhibited most major DSB repair factors from both the NHEJ and HR pathways, characterising their mutational profiles in unprecedented detail.

## RESULTS

### Overview of the iMUT-seq technique

To provide controlled induction of DSBs at known genomic loci, we employed the damage-induced via AsiSI (DIvA) cell-system. Created by the Legube Lab, DIvA utilises an AsiSI restriction enzyme fused to an oestrogen receptor to allow for 4-hydroxytamoxifen (OHT) inducible DSBs at the AsiSI recognition sites across the human genome. In particular, we chose the auxin-inducible degron version of this system (AID-DIvA), that allows for the subsequent degradation of the AsiSI enzyme via treatment with indole-3-acetic acid (IAA), therefore preventing further DSB induction and allowing the cells to repair the current DSBs (Figure 1A,B,C). Previous research has profiled the AsiSI recognition sites that are reproducibly cut by the AsiSI-ER fusion protein (Clouaire *et al*, 2018) and also characterised subsets of these that are preferentially targeted for either NHEJ or HR repair (Aymard, F. *et al*, 2014).

**Figure 1:**
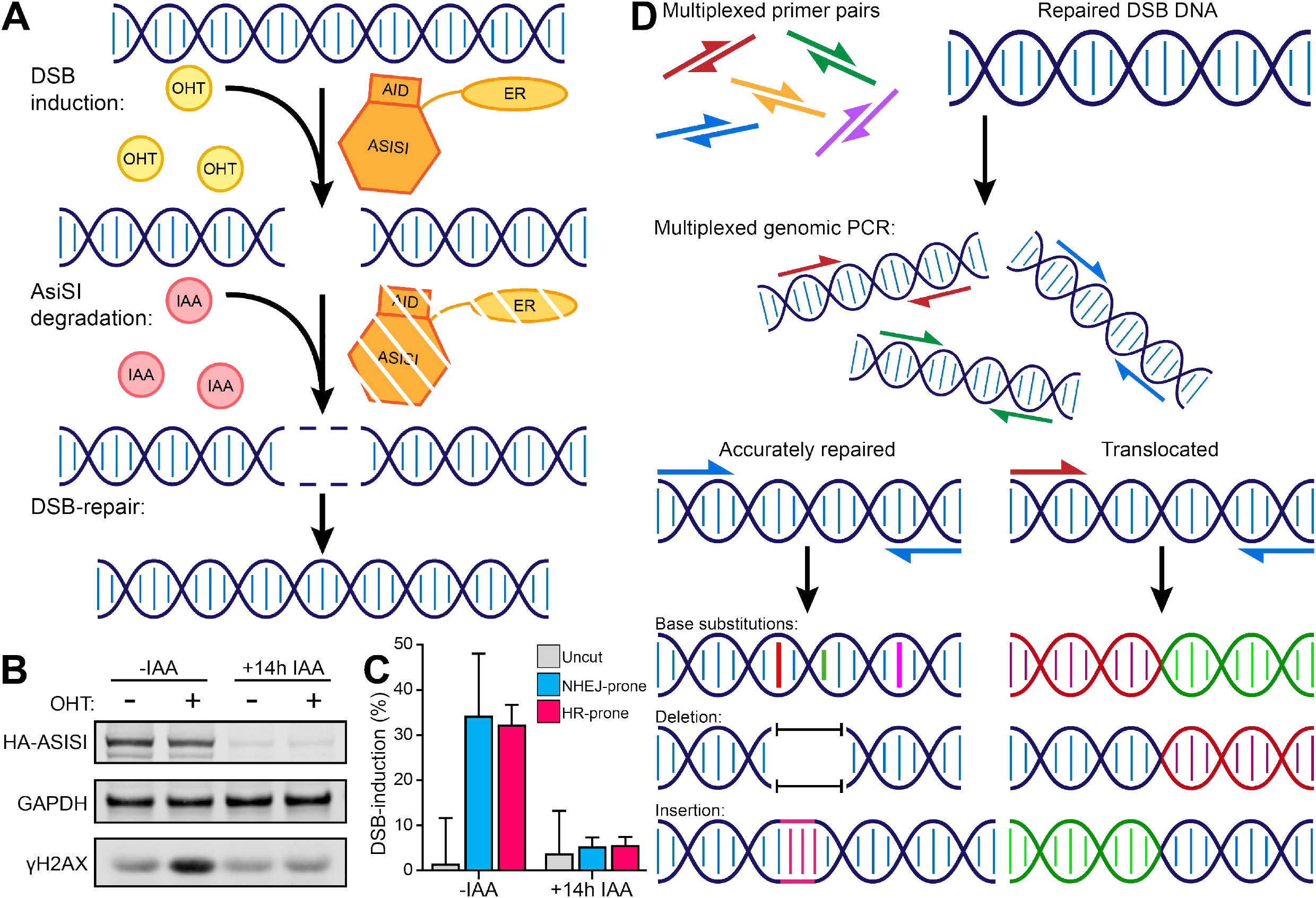
Overview of the iMUT-seq technique. **(A)** Experimental design of iMUT-seq, first Damage-Induced via AsiSI (DIvA) cells are treated with hydroxytamoxifen (OHT) to translocate the AsiSI-ER-AID fusion protein to the nucleus where it generates DSBs by cutting it’s recognition sequences spread across the genome. Next, the fusion protein is degraded by activating it’s auxin-inducible degron (AID) with auxin (IAA) treatment, allowing the cell to fully repair all the AsiSI induced DSBs. **(B)** iMUT-seq sequencing pipeline. First, genomic DNA from (A) is amplified in a PCR reaction with a multiplex of primers targeting the AsiSI induced DSB sites. This amplifies both correctly repaired DSBs as well as translocated DSBs, since all the primers are in the same PCR reaction, and therefore primers from different pairs can amplify the translocated DNA. This amplified DNA can then be sequenced via NGS, allowing us to profile mutations around DSBs at single-nucleotide resolution, as well as map translocations across the genome. **(C)** Western blot of AID-DIvA cells treated with OHT to induce DSBs, demonstrated by increased γH2AX, and IAA to induce degradation of the AsiSI fusion protein, with GAPDH as a loading control. **(D)** Representation of qPCR quantification of DSB induction at 3 different loci amplified in iMUT-seq; one NHEJ-prone locus, one HR-prone locus and one uncut control locus. DSB induction is calculated as a delta Ct normalised to without OHT treatment.

We then coupled this approach to a multiplexed genomic PCR that amplifies 25 regions: 10 NHEJ-prone, 10 HR-prone and 5 uncut control loci (Table S1), followed by NGS to sequence over repaired, endogenous DSBs at very high depth (Figure 1D), accurately quantifying mutations at rates as low as 1 in 200,000 events. The initial genomic PCR products are ∼250-290bp in length (Figure S1A) so that paired-end 150bp sequencing can give full coverage of the entire amplicon, which excluding the primer sequences extend at least 100bp either side of these DSBs. These amplicons can then be used in any standard DNA library preparation protocol to create libraries ready for NGS (Figure S1A).

In addition, translocation events are known to occur between DSBs (Richardson and Jasin, 2000; Aymard, François *et al*, 2017) and are known to be critical in cancer biology (Nowell and Hungerford, 1961; Richardson and Jasin, 2000), therefore their quantification is an important component of investigating DSB-induced mutagenesis. With all primers in a single PCR reaction, translocated loci will still be amplified by the corresponding upstream and downstream primers and can be subsequently quantified within the sequencing results (Figure 1D).

We noted that DSB-induced samples had significantly reduced alignment efficiency due to their mutations. To address this, we used a machine learning approach that utilises a genetic algorithm to systematically test different alignment parameters, optimising the alignment efficiency over sequential rounds of optimisation and re-testing (Figure S1B). The optimal alignment parameters identified by this significantly increased alignment efficiency, reducing the number of unaligned reads in our results and removing the difference between undamaged and damaged samples (Figure S1C).

### iMUT-seq profiling of HR and NHEJ-dependent DSB induced mutations

Previous investigations into DSB-induced mutations have been conducted, primarily focusing on deletion and insertion events at breaks (Ahrabi *et al*, 2016; Hussmann *et al*, 2021; Schep *et al*, 2021). However, no studies have fully mapped the mutations around DSBs, especially at endogenous loci, and far more detail is required to understand pathway specific mutations at DSBs. To fully investigate the mutations that occur around endogenous DSBs and to explore the differences in mutagenesis between the major DSB repair pathways, we initially compared the mutation profiles of DSB loci that are prone to either NHEJ or HR repair.

We first profiled base substitutions which found a strong peak of mutations that spread to ∼25bp either side of the DSB (Figure 2A). Interestingly, HR-prone loci show a clear increase in mutation rates compared to NHEJ-prone loci (Figure 2B) with NHEJ-prone loci also having a narrower peak around the break, only spreading to ∼10bp either side of the DSB (Figure 2A). This is despite previous analysis showing that both HR and NHEJ-prone sites have similar rates of cutting by the AsiSI enzyme (Aymard, François *et al*, 2017). Analysing the base substitution signatures found that C>T mutations are most common, closely followed by C>G and C>A mutations, and subsequently T>C, T>G and T>A mutations respectively (Figure 2C). This signature is most closely aligned to COSMIC signature 3 (Bamford *et al*, 2004) which is well characterised as a DSB-induced mutation signature commonly found in breast, ovarian and pancreatic cancers. Interestingly, there are also similarities with COSMIC signature 5, which has an unknown aetiology.

**Figure 2:**
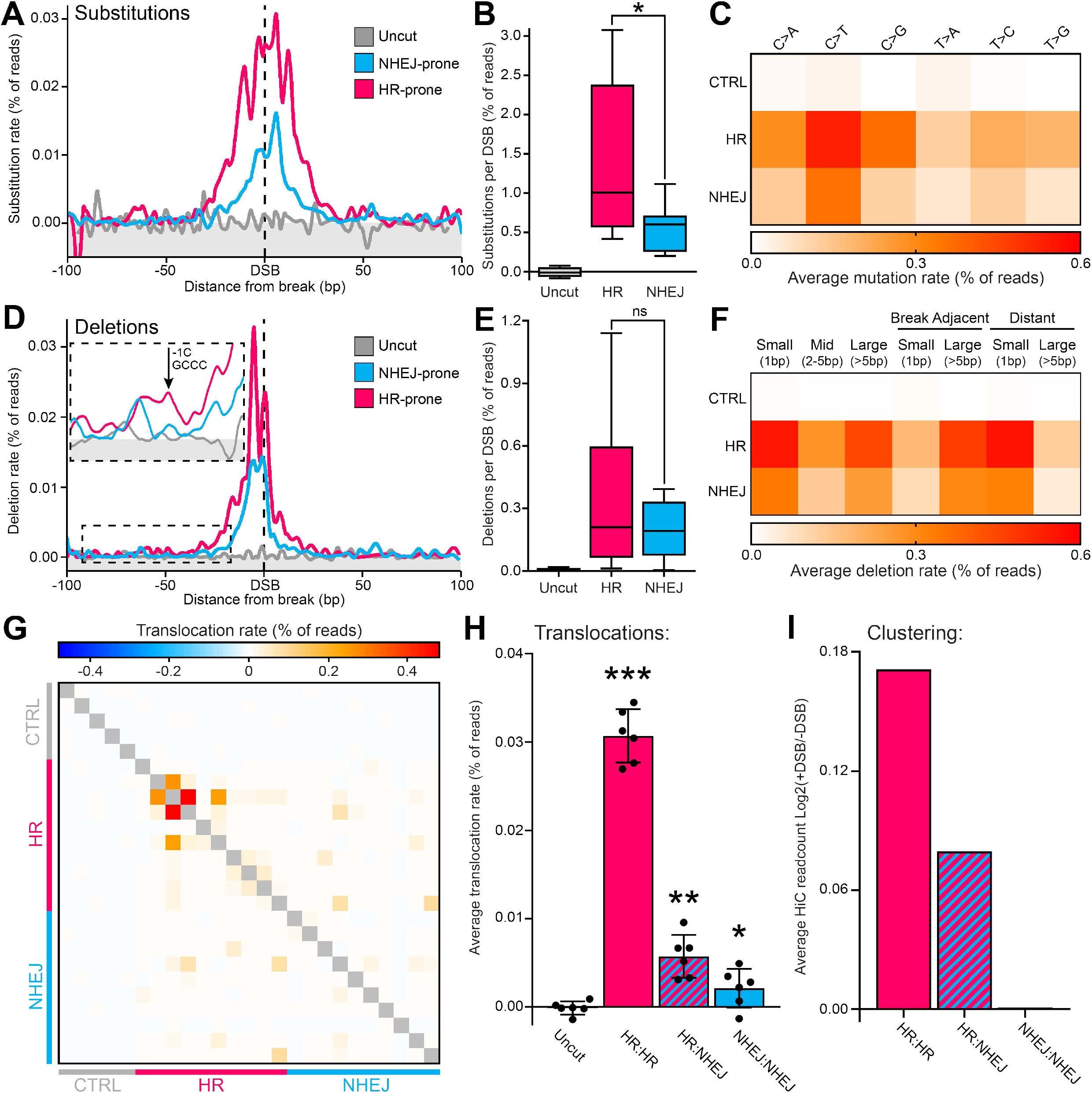
iMUT-seq profiling of HR and NHEJ-dependent DSB induced mutations. **(A)** Metagene line plot of base substitution rate quantified by iMUT-seq, as a percentage of readcount, 100bp either side of AsiSI induced DSBs prone to repair by either NHEJ or HR, or at uncut control loci. **(B)** Boxplot of total base substitutions per loci at either uncut control, HR-prone or NHEJ-prone loci quantified by iMUT-seq, statistics done via unpaired Wilcoxon test, * p<0.05. **(C)** Heatmap of the average rate of each base substitution type per DSB loci at either uncut control, HR-prone or NHEJ-prone loci, quantified by iMUT-seq. **(D)** Same as (A) but for deletions, also with a zoomed in section in the upper left showing deletions at distance from the break point and highlighting the mononucleotide deletions caused by polymerase slippage at polynucleotide repeats. **(E)** Same as (B) but for deletions. **(F)** Heatmap of average rate of different types of deletion per loci at either uncut control, HR-prone or NHEJ-prone loci, quantified by iMUT-seq. **(G)** Heatmap of translocations quantified by iMUT-seq, each row and each column represent different iMUT-seq amplicons of either uncut control, HR-prone or NHEJ prone loci with each cell being the translocation rate as a percentage of total readcount delta to -DSB between the two loci that correspond to that cell. **(H)** The average translocation rate between different loci quantified by iMUT-seq, points represent each biological replicate and error bars are S.D., all statistics are done relative to the uncut control results using a paired t-test, * p<0.05, ** p<0.01, *** p<0.001. **(I)** Average clustering between the different DSB loci in (H), determined by log2 +DSB/-DSB in Capture-HiC (Aymard, François *et al*, 2017), showing that DSB clustering is very similar to translocation rates shown in (H).

Quantification of deletions found a similar peak of mutations centred on the break, although this peak was more concise than for base substitutions (Figure 2D). Deletions only showed a subtle and insignificant difference in rate between HR and NHEJ prone sites (Figure 2E), which suggests a relatively higher propensity for NHEJ repaired sites to undergo deletions compared to base substitutions. Investigating deletion length found that small deletions (mononucleotide, 1bp) were the most common, although larger deletions, classified as either mid (2-5bp) or large (>5bp) length, also occurred relatively frequently (Figure 2F). Large deletions are almost all adjacent to the break point, whereas a majority of small deletions are spread at distance from the break point (Figure 2F, S2A). Further investigation of these distant deletions revealed that they commonly occur at polynucleotide repeats (Figure 2D), suggesting that they are caused by polymerase slippage when filling in resected overhangs (McVey *et al*, 2016; Guirouilh-Barbat *et al*, 2014; Yang *et al*, 2008; Hicks, Kim and Haber, 2010; Deem *et al*, 2011). We also quantified the rate of insertion mutations, however this yielded very few DSB-induced mutations, especially compared to deletion rates (Figure S2B-C), with no statistically significant difference between uncut and DSB induced loci (Figure S2C). This indicates that although possible, insertions occur relatively infrequently at DSBs compared to other mutation types, which is consistent with previous observations when directly comparing the frequency of deletions and insertions (Ahrabi *et al*, 2016; Song *et al*, 2021).

Repair of DSBs using alternative end-joining (aEJ) pathways is commonly thought to occur as a backup process in case of failed HR or NHEJ. This is commonly thought to occur through microhomology-mediated end-joining (MMEJ) which utilises short stretches of homology, created by digestion of the broken DNA ends, to facilitate the ligation of these sticky ends (Wang and Xu, 2017; Decottignies, 2013; Bennardo *et al*, 2008). We are therefore able to quantify these microhomologies, which although occur very infrequently compared to regular deletions (Figure S2D-E) do represent larger than average deletions (Figure S2F). Despite these deletions stretching up to ∼70bp, all of the homologous stretches were 2-7bp in length which is known to be common for MMEJ (Figure S2G) (Ahrabi *et al*, 2016; Schep *et al*, 2021; Decottignies, 2013; Wang and Xu, 2017).

Translocation mapping yielded an interesting bias in the frequency of translocations towards particular loci (Figure 2G). Specifically, translocation events between HR loci and other HR loci are a majority of the translocation events (Figure 2H) and this is largely due to a few specific loci (Figure 2G). Previous reports have demonstrated a mechanism of clustering of active DSBs within the nucleus, where DSBs at different loci spatially group together within the nucleus in response to break formation (Aymard, François *et al*, 2017). This mechanism was suggested to be a source of translocations, and we therefore analysed the clustering of the loci we sequenced, which found that clustering of these loci is very comparable to their translocation frequencies (Figure 2I), suggesting that translocation at DSBs is driven by this mechanism.

To statistically determine the sensitivity of iMUT-seq, we calculated for each nucleotide around the DSB the statistical significance of the mutation induction at DSB loci compared to the uncut control loci (Figure S2I). This revealed that iMUT-seq can consistently identify mutations that occur as infrequently as 0.005%, or 1 in 200,000 (Figure S2I). Although some rarer mutations were also identified, the low induction and increased variability resulted in these mutation calls being uncommon and only borderline significant (Figure S2J). We are therefore confident that iMUT-seq can routinely identify mutations as rare as 1 in 200,000 events, although it is possible that this could be increased by increasing read depth.

On average, we found DSBs to have a mutation rate per site of ∼1% for base substitutions, ∼0.25% for deletions and ∼0.01% for translocations, while insertions have little to no significant induction (Figure 2B,E, S2H, S2C), though these numbers are likely an underrepresentation due to the frequency of AsiSI cutting being only ∼25-30% per loci (Aymard, François *et al*, 2017; Cohen *et al*, 2018; Lu *et al*, 2018). From these results, we can also infer that NHEJ is more prone to deletions at the break point, whereas HR is more prone to mutations from polymerase error and slippage due to re-polymerisation of resected DNA during HR. We also see a distinct bias in the frequency of both base substitutions and translocations towards HR repaired loci; however, due to the cell-cycle dependency of HR, NHEJ will frequently repair HR-prone loci. It is therefore possible that HR-prone loci have a higher propensity for mutations, which could also contribute to why they are prone to the higher-fidelity HR pathway. This is further explored as we examine the effects of depletion of NHEJ and HR factors and their influence on these mutations.

### Disrupting late stage HR or NHEJ factors results in increased mutation rates compared to initiating factors

To fully investigate the mutagenic mechanism of NHEJ and HR repair, we conducted siRNA mediated depletion of the following NHEJ factors: KU70, Artemis, 53BP1, Pol-λ, XRCC4 and LIG4 and HR factors: MRE11, BRCA1, BLM, EXO1, BRCA2, FANCA, RAD52, RAD51, Pol-δ and Pol-ε (Table S2) (Figure S3A), as well as the chemical inhibition of DNA-PK and ATM.

An overall analysis of the impact of these treatments found that there are distinct trends in DSB-induced mutations as you interfere with different stages of both NHEJ and HR. Depletion of early repair pathway components resulted in a reduction of all types of mutations relative to control siRNA, whereas depletion of components further down the repair pathways resulted in a progressive increase in mutagenesis, ultimately leading to significantly increased mutations when the late NHEJ factor LIG4 and the late HR factor RAD51 were depleted (Figure 3A). There are notable exceptions to this trend, such as DNA-PKi treatment which caused a remarkable increase in base substitutions and large deletions, likely due to the specific mechanism of DNA-PK kinase inhibition (DNA-PKi) (Uematsu *et al*, 2007). Depletion of the HR polymerases Pol-δ and Pol-ε also did not follow this trend, resulting in only small changes in mutations likely due to a level of redundancy in the various polymerases that have been found to function during HR repair (McVey *et al*, 2016).

**Figure 3:**
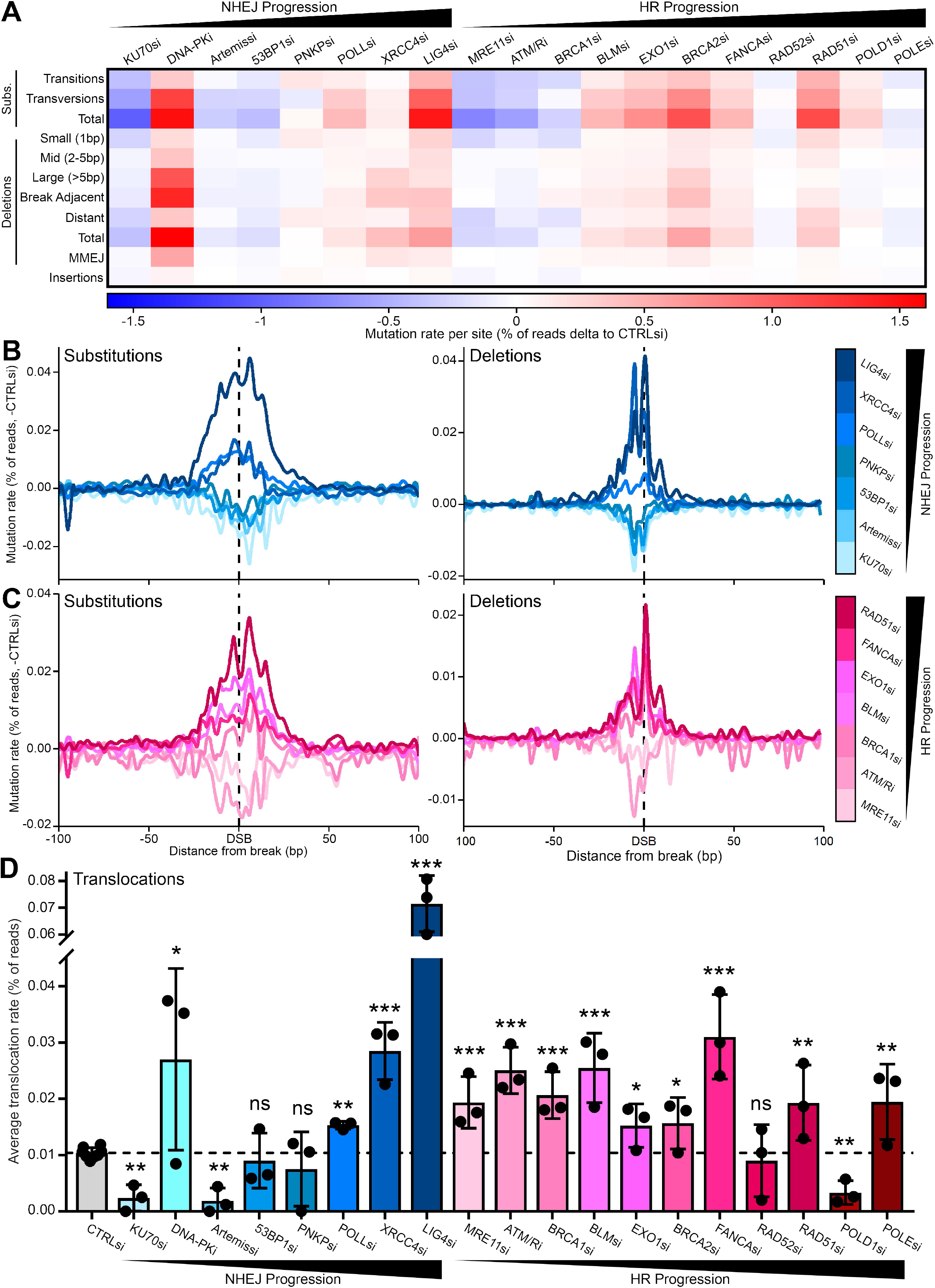
Interrupting DSB repair causes increasing mutation rates the later the interruption occurs in the pathway. **(A)** Heatmap of the average rate per DSB locus of different mutation types, calculated as a percentage of readcount delta to -DSB then delta to control siRNA for 19 different DDR targeting siRNAs, quantified by iMUT-seq. **(B)** Metagene line plots of base substitutions rate (left) and deletion rate (right) as a percentage of readcount delta to -DSB then delta to control siRNA, 100bp either side of AsiSI induced DSBs upon knockdown of several different NHEJ repair factors, quantified by iMUT-seq. Legend (right) shows the colours for each siRNA target, as well as a bar depicting the position of the factors in the progression of NHEJ repair. **(C)** Same as (B) but with the knockdown of several HR repair factors. **(D)** Average translocation rate between DSBs quantified by iMUT-seq as a percentage of readcount upon knockdown of 19 different DSB repair factors from both NHEJ and HR repair points represent each biological replicate and error bars are S.D.,, all statistics are done relative to the control siRNA result using a paired t-test, * p<0.05, ** p<0.01, *** p<0.001.

We next investigated this mutation trend spatially across break sites. This similarly found that DSB induced mutations increase the later the defect occurs in either NHEJ (Figure 3B) and HR (Figure 3B-C) pathways. We found that mutations proportionally change across the break sites, with mutations often spreading further from the break as their frequency increased (Figure 3B-C). Analysing how these treatments alter deletion lengths also found this trend, with early NHEJ factor depletion reducing deletion length whereas late NHEJ factor depletion significantly increased deletion length (Figure S3B). Although early HR factor depletion did not reduce deletion length, depletion of later factors significantly increased deletion length with Pol-δ having the strongest increase of all HR factors (Figure S3B).

Finally, investigation of translocation rates found that depletion of NHEJ factors continued to follow this trend (Figure 3D), however HR factors did not. Depletion of early HR factors significantly increased translocation rates and did so to a similar degree of later HR factors (Figure 3D). This suggests that prevention of translocations is a key function of HR, more so than the prevention of base substitutions or deletions since these decrease with early inhibition of HR (Figure 3A-C).

This overall analysis has highlighted trends in the mutation profile of DSB repair factor depletion. The overriding trend shows that depletion of repair factors has a greater impact on mutagenesis the later into the pathway that the repair factor acts. This resulted in groups of siRNA targets that can loosely be defined as early NHEJ, late NHEJ, early HR and late HR repair, though with some clear exceptions to this rule, such as DNA-PKi due to it’s mechanism of action (Uematsu *et al*, 2007; Reginato, Cannavo and Cejka, 2017) and the HR polymerases Pol-δ and Pol-ε which likely have redundant roles (McVey *et al*, 2016). In addition, loss of either KU70 and MRE11, the initiating components of NHEJ and HR respectively, resulted in similar reductions in base substitutions and deletions, suggesting that both NHEJ and HR contribute to the mutation profiles observed previously (Figure 2).

### NHEJ induces mutations around DSBs but protects against large-scale loss of genome integrity

The early NHEJ factors KU70, Artemis, 53BP1 and PNKP all showed a similar mutation profile in our initial analysis (Figure 3). Further investigation of these factors showed a clear trend as they all significantly reduced base substitutions and deletions around the break and all have very comparable profiles (Figure 4A-B). KU70 depletion also exhibited a shift in base substitution signature, altering these mutations towards T>C and T>G mutations while reducing all others, despite the other early NHEJ factors not having a significant effect (Figure 4C). Interestingly, KU70 depletion was also distinct from other factors at the deletion level. Knockdown of KU70 significantly reduced all types of deletion, however knockdown of Artemis, 53BP1 and PNKP all reduced medium and especially large, break adjacent deletions, but caused an increase in small distant deletions (Figure 4D). This is consistent with these factors suppressing HR, as their depletion would cause a switch towards the HR signature of increased distant, mononucleotide deletions due to resection of the DNA leading to increased polymerase errors, as well as a reduction in the NHEJ signature of large break-adjacent deletions. However, it is unusual that KU70 depletion does not also follow this trend, it could be due to reduced NHEJ signalling leading to a loss of competition with NHEJ processes and thus reduced HR-induced mutations.

**Figure 4:**
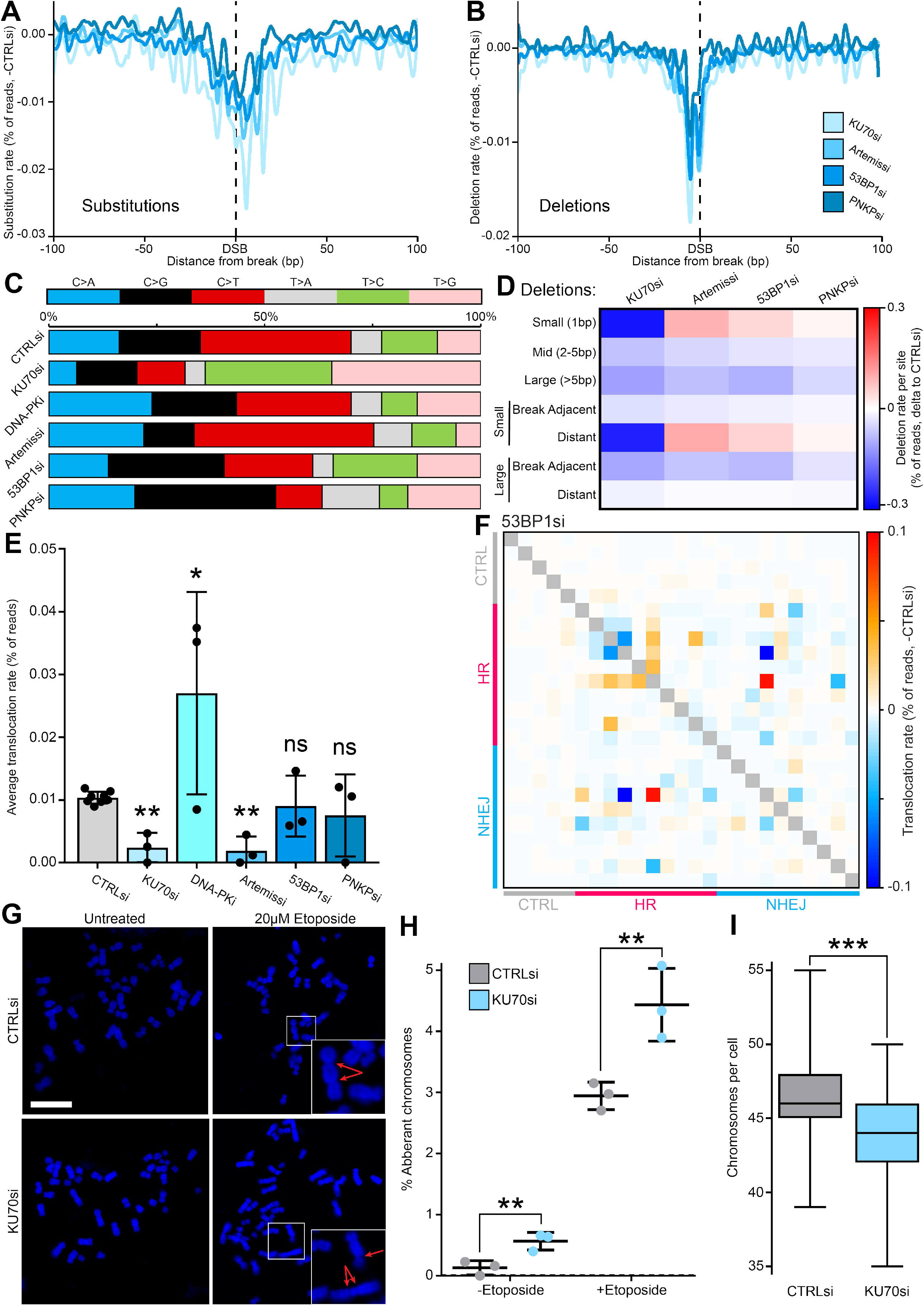
NHEJ induces mutations around DSBs but protects against large-scale loss of genome integrity. **(A)** Metagene line plot of base substitution rate as a percentage of readcount delta to -DSB then delta to control siRNA, 100bp either side of AsiSI induced DSBs upon knockdown of several different NHEJ repair factors, quantified by iMUT-seq. **(B)** Same as (A) but for deletion rate. **(C)** Stacked bar plot of the relative base substitution rates per DSB locus quantified by iMUT-seq upon depletion or inhibition of different early NHEJ factors. **(D)** Heatmap of the average rate per DSB loci of different deletion types, calculated as a percentage of readcount delta to -DSB then delta to control siRNA, quantified by iMUT-seq. **(E)** Average translocation rate between DSBs quantified by iMUT-seq as a percentage of readcount delta to -DSB, points represent each biological replicate and error bars are S.D., all statistics are done relative to the control siRNA result using a paired t-test, * p<0.05, ** p<0.01. **(F)** Heatmap of translocation rates between different DSBs quantified by iMUT-seq with 53BP1 depletion, each row and each column represent different iMUT-seq amplicons of either uncut control, HR-prone or NHEJ prone loci with each cell being the translocation rate as a percentage of total readcount delta to -DSB between the two loci that correspond to that cell and then delta to control siRNA. **(G)** Representative images of metaphase spreads in HCT116 cells with or without 20μM etoposide treatment and with either control or KU70 siRNA mediated depletion, scalebar is 10µm. Zoom in sections highlight chromosomal rearrangements, single arrows indicate chromosome breaks, double arrows indicate dicentric chromosomes. **(H)** Quantification of chromosomal aberrations in the metaphase spreads from (G) as a percentage of total chromosome number, points represent each biological replicate and error bars are S.D., statistics done relative to control siRNA using a paired t-test, ** p<0.01. **(I)** Quantification of chromosome number per cell in metaphase spreads from (G), statistics done using an unpaired Wilcoxon test, *** p <0.001.

Treatment with an inhibitor of the DNA-PK kinase yielded remarkably differing results from these siRNA mediated depletions. DNA-PKi treatment results in strong induction of both base substitutions and deletions specifically at the break point (Figure S4A). This showed the strongest mutation induction of all of our treatments (Figure 3A); however, this is skewed towards break adjacent deletions (Figure S4A, Figure 3A). Studying the lengths of these DNA-PKi induced deletions showed that inhibition of DNA-PK significantly increases all deletion lengths, but was heavily skewed towards increased large deletions (Figure S4B). This mutation signature greatly supports the mechanism of DNA-PKi locking the DNA-PK complex onto DNA ends due to the loss of its autophosphorylation activity (Uematsu *et al*, 2007), leading to the endonucleolytic cleavage of the DNA-PK bound DNA-ends by the MRN complex (Shibata *et al*, 2014; Reginato, Cannavo and Cejka, 2017), as this would specifically produce large deletions at the break site.

Studying the effect of these early NHEJ disruptions on translocations gave similar results to other mutations, with all depletions reducing or not significantly altering translocation rates (Figure 4E), except for DNA-PKi which results in an increase (Figure 4E, S4C). 53BP1 depletion did not significantly decrease translocation rates, but a look at the mapping of these translocations revealed that this is actually due to some events decreasing in frequency while others increase (Figure 4F). This suggests that early NHEJ could have a role in suppressing translocations at certain genomic loci or under certain conditions.

The depletion of early NHEJ factors causing reduced mutations does support the notion of NHEJ being a mutagenic repair process, and the especially the reduction in translocations seen with KU70 and Artemis depletion as genomic rearrangements are thought to commonly be the result NHEJ (Ghezraoui *et al*, 2014; Cannan and Pederson, 2016). However, this raises the question of the function of NHEJ in DSB repair as a whole. To gain further insight into this, we conducted metaphase spreads with etoposide treatment in HCT116 cells to study the effects of KU70 depletion on a genomic scale as an orthogonal validation. Remarkably, this showed that KU70 depletion significantly increases chromosomal aberrations (Figure 4G-H), implicating NHEJ in suppressing genomic rearrangements. Quantifying chromosome numbers per cell revealed that KU70 depletion also results in a significant loss of chromosomes in response to etoposide treatment (Figure 4I). These results indicate that NHEJ is necessary for genome maintenance, specifically via the prevention of large-scale chromosomal aberrations and DNA loss. We therefore believe that in the absence of NHEJ, repair is significantly delayed resulting in substantial deletion of the DNA at the break and causing either mutagenic repair, by pathways such as MMEJ or single-strand annealing (SSA), or DNA loss via mechanisms such as micronuclei formation. These large-scale genomic aberrations would likely not be detectable by iMUT-seq as they would result in the deletion of the primer regions used for the genomic PCR, leading to a perceived reduction in translocations.

Collectively, these results highlight a key role for NHEJ in maintaining large-scale genome integrity and the suppression of HR induced distant mutations. In addition, we found a crucial caveat in our iMUT-seq technique where treatments that induce substantial loss of genome integrity lead to inaccurate results which are captured by metaphase spread analysis. Since metaphase spreads are an effective and well-established approach for the quantification of chromosomal rearrangements, and they provide an overall view of the genome they make an excellent validation technique to pair with iMUT-seq which interrogates highly specific mutations on a nucleotide level.

### Disruption of late NHEJ processes promotes MMEJ deletions and translocations

The knockdowns of POLL, XRCC4 and LIG4 showed a significant deviation from the early NHEJ phenotype in our overall analysis (Figure 3). A closer look revealed that all three of these knockdowns result in increased base substitutions and deletions around break sites, although they present with very different profiles (Figure 5A-C). There were no notable alterations in base substitutions signatures between these depletions (Figure S5A).

**Figure 5:**
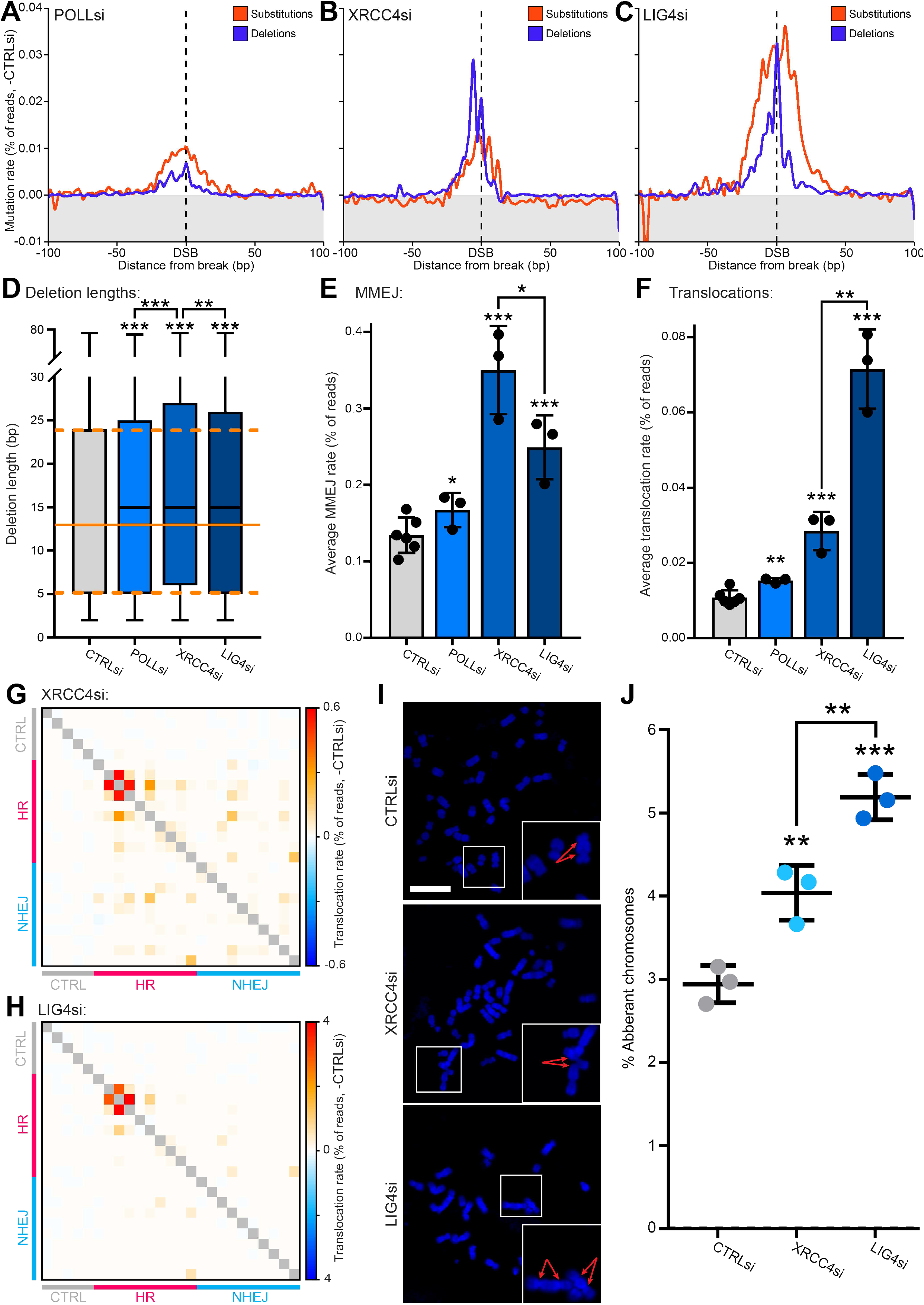
Disruption of late NHEJ processes promotes MMEJ deletions and translocations. **(A)** Metagene line plot of base substitution and deletion rates as a percentage of readcount delta to -DSB and then delta of POLL siRNA – control siRNA, 100bp either side of AsiSI induced DSBs, quantified by iMUT-seq. **(B)** Same as (A) but for XRCC4 siRNA. **(C)** Same as (A) but for LIG4 siRNA. **(D)** Boxplot of DSB-induced deletion lengths quantified by iMUT-seq upon knockdown of POLL, XRCC4 or LIG4, all statistics are done relative to the control siRNA result using an unpaired Wilcoxon test, *** p<0.001. **(E)** Average rate of microhomology-mediated end-joining (MMEJ) per DSB loci quantified by iMUT-seq as a percentage of readcount delta to -DSB, points represent each biological replicate and error bars are S.D., all statistics are done relative to the control siRNA result using a paired t-test, * p<0.05, ** p<0.01, *** p<0.001. **(F)** Same as (E) but for translocations per DSB locus. **(G)** Heatmap of translocation quantified by iMUT-seq with XRCC4 depletion, each row and each column represent different iMUT-seq amplicons of either uncut control, HR-prone or NHEJ prone loci with each cell being the translocation rate as a percentage of total readcount delta to -DSB between the two loci that correspond to that cell and then delta to control siRNA. **(H)** Same as (G) but for LIG4 siRNA. **(I)** Representative images of metaphase spreads in HCT116 cells with 20μM etoposide treatment and with either control, XRCC4 or LIG4 siRNA mediated depletion, scalebar is 10µm. Zoom in sections highlight chromosomal rearrangements, double arrows indicate dicentric chromosomes. **(J)** Quantification of chromosomal aberrations in the metaphase spreads from (I) as a percentage of total chromosome number, points represent each biological replicate and error bars are S.D., statistics done relative to control siRNA using a paired t-test, ** p<0.01, *** p<0.001.

However, further interrogation of the POLL base substitution profile did show an interesting difference between NHEJ and HR-prone loci. POLL depletion led to an increase in distant substitutions specifically at HR-prone loci (Figure S5B). This implicates POLL in gap-filling at sites that have undergone short-range resection during early HR repair, suggesting a switch from HR to NHEJ.

Comparing deletion lengths revealed that depletion of these factors results in a general increase in the length of deletions at breaks, with XRCC4 having the strongest effect (Figure 5D). By breaking down deletion frequency by length of deletion, we see that whereas LIG4 depletion has the greatest overall increase in deletions (Figure 3A, 5C), this is due to an increase in small, mid and large deletions, whereas POLL and XRCC4 specifically increase mid and large deletions (Figure S5C). In particular, XRCC4 knockdown substantially increased large deletions, even showing a significant increase over LIG4 knockdown (Figure S5C). Examining MMEJ revealed that this is due to XRCC4 depletion leading to a considerable increase in the rate of MMEJ, even relative to LIG4 and POLL which also see increases in MMEJ (Figure 5E). This disparity in mutagenic mechanism is particularly interesting given the close mechanistic relationship between XRCC4 and LIG4 in the repair pathway (Conlin *et al*, 2017; Andres *et al*, 2012; Roy *et al*, 2012).

Comparing the frequencies of translocations between these knockdowns found that although all result in increased translocations, LIG4 depletion results in an extraordinary increase in translocations (Figure 5F). Both XRCC4 and LIG4 depletion show similar translocation maps (Figure 5G-H), with both resulting in a global increase in translocations across different loci (Figure S5D). Thus, while XRCC4 depletion promotes MMEJ, LIG4 depletion strongly promotes translocation between DSBs. To validate this differential in translocation rates, we again conducted metaphase spreads in HCT116 cells using etoposide treatment in combination with XRCC4 and LIG4 knockdown. This experiment confirmed our iMUT-seq results, showing a significant increase in chromosomal aberrations with both XRCC4 and LIG4 depletion, but with LIG4 also displaying an increase over XRCC4 (Figure 5I-J).

These results clearly demonstrate significant delineation in the mutation signatures of later NHEJ repair factors. This is likely due to the different roles these proteins have in factor recruitment and DNA processing leading to profound impacts on the mutagenic mechanisms that occur when they are depleted.

### HR repair induces deletions and base substitutions to prevent translocations

In studying the role of early HR factors, we utilised small molecule inhibitors of ATM (10μM KU55933) and ATR (10μM VE-821) kinases. Initially we conducted separate and combined inhibition of the kinases to determine how redundant their functions are for these repair processes. Separate ATM and ATR inhibition lead to similar phenotypes, however this signature is significantly increased with combination treatment (Figure S6A). This supports the theory of redundancy in these kinases during DSB repair (Blackford and Jackson, 2017), and we therefore used the combination treatment of ATM and ATR inhibitors in our analysis.

Initial analysis showed a decrease in mutations seen when depleted early HR factors (Figure 3A-C), as HR is generally considered a high-fidelity pathway that prevents such mutations. A specific investigation of these factors found that MRE11 depletion and ATM/ATR inhibition both reduce base substitutions and deletions across the entire break, however BRCA1 depletion promotes mutations specifically adjacent to the break, while reducing distant mutations (Figure 6A-B). Base substitution signatures showed that MRE11 depletion promotes C>G, but reduces C>A mutations, while BRCA1 depletion promotes T>A, but reduces C>T mutations (Figure S6B), which could be indicative of the mechanisms by which repair progresses when these factors are depleted. Interrogation of deletion rates found that all these treatments resulted in decreased distant mononucleotide deletions, while BRCA1 and, to a lesser extent, MRE11 depletions specifically reduced large break adjacent deletions, but increased small and mid-deletions at the break (Figure 6C, S6C-E). This shows a shortening of the deletions upon prevention of early HR, suggesting that short-range resection promotes longer deletions at DSBs. Although ATM/ATR inhibition did not lead to an increase in smaller break-adjacent deletions the reduction in deletions still skewed towards large deletions (Figure 6C).

**Figure 6:**
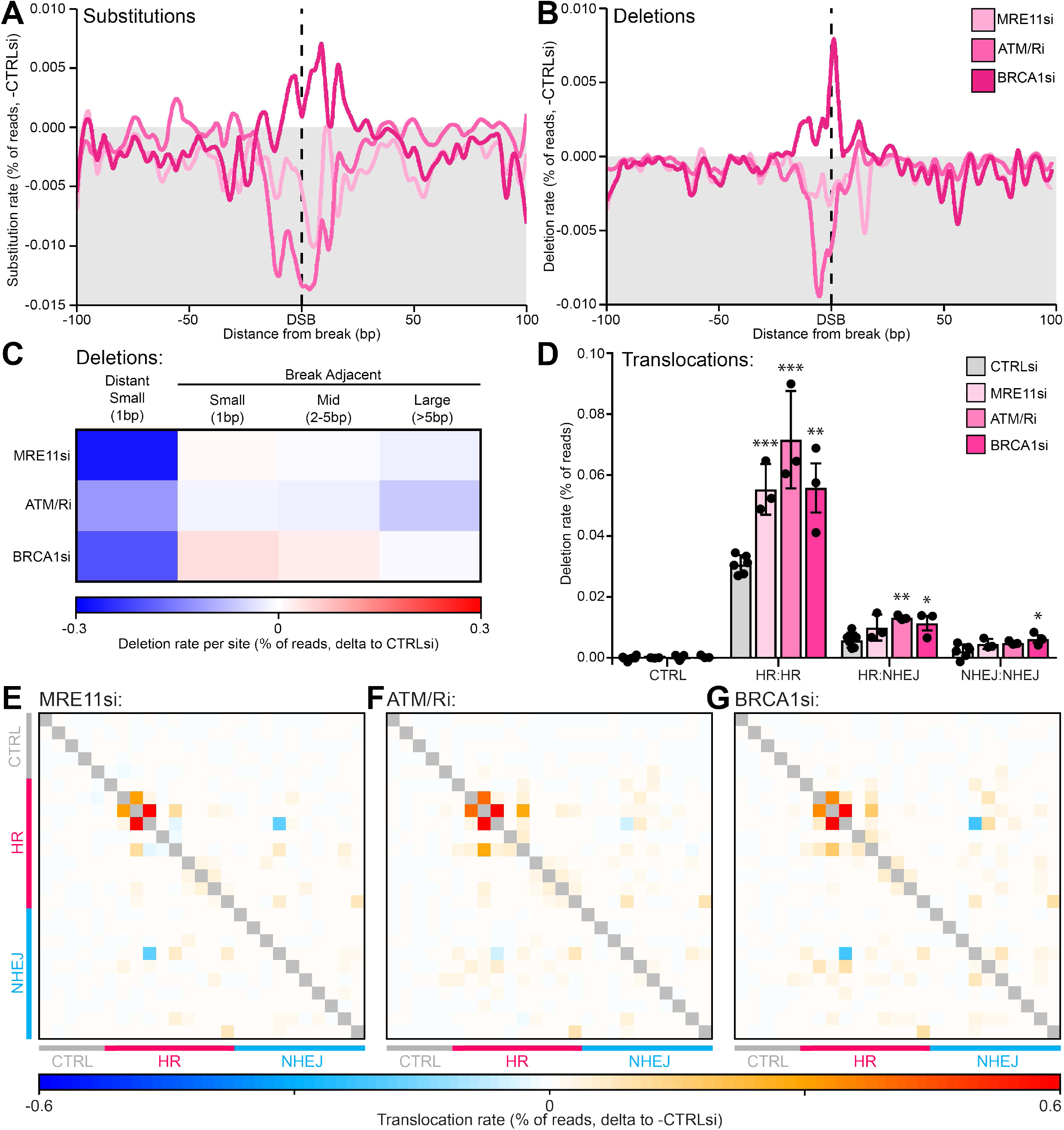
Initiation of DSB resection increases mutations and deletion length but suppresses translocations. **(A)** Metagene line plot of base substitution rate as a percentage of readcount delta to -DSB then delta to control siRNA/mock treatment, 100bp either side of AsiSI induced DSBs upon knockdown or inhibition of MRE11, ATM/ATR or BRCA1, quantified by iMUT-seq. **(B)** Same as (A) but for deletion rate. **(C)** Heatmap of the average rate per DSB loci of different deletion types, calculated as a percentage of readcount delta to -DSB then delta to control siRNA, quantified by iMUT-seq. **(D)** Average translocation rate between different DSBs quantified by iMUT-seq, split by events between sites repaired by either HR or NHEJ or uncut control loci, with depletion or inhibition of MRE11, ATM/ATR or BRCA1, points represent each biological replicate and error bars are S.D., all statistics done relative to control siRNA using paired t-tests, * p<0.05, ** p<0.01, *** p<0.001. **(E)** Heatmap of translocation rates quantified by iMUT-seq with MRE11 depletion, each row and each column represent different iMUT-seq amplicons of either uncut control, HR-prone or NHEJ prone loci with each cell being the translocation rate as a percentage of total readcount delta to -DSB between the two loci that correspond to that cell and then delta to control siRNA. **(F)** Same as (E) but with ATM/ATR inhibition. **(G)** Same as (E) but with BRCA1 depletion.

Analysis of translocation rates shed further light on the mutagenic mechanisms of early HR, as all treatments significantly increased translocations (Figure 6D). All treatments showed similar translocation maps with generally increased frequencies (Figure 6E-G), suggesting similar mechanistic impacts on translocations, with BRCA1 even inducing translocations between NHEJ-prone sites (Figure 6D).

These results, in conjunction with the data presented earlier (Figure 3), question the role of HR in preventing nucleotide level mutations, such as substitutions and deletions. Instead, they suggest that HR promotes these mutations as resection leads to polymerase errors and slippage as a result of re-polymerisation of the resected DNA. In addition, we found that loss of resection shortens the deletions that occur at break ends, indicating that the resected DNA overhang is susceptible to degradation and therefore resection promotes larger deletions at DSBs.

### BLM and EXO1 synergistically protect against different mutations signatures

Long-range resection is a defining step in HR and is thought to be promoted by a variety of enzymes, with EXO1 and the BLM-DNA2 complex chief among these. A comprehensive analysis of how BLM and EXO1 depletion influence mutation signatures is therefore necessary to characterise any mutagenic differences in repair involving these enzymes.

BLM and EXO1 knockdown present very similar mutation profiles, exhibiting a strong peak of base substitutions and deletions around the break, although with EXO1 depletion showing a clearly higher level of mutations than BLM depletion (Figure 7A-B). This feature was seen across all mutation types, with both BLM and EXO1 knockdown increasing mutations compared to control, but EXO1 knockdown also showing an increase compared to BLM (Figure 7C). Both treatments also showed remarkably similar base substitution signatures, skewing towards C>G and away from C>T mutations (Figure S7A). This suggests that both BLM and EXO1 knockdown share mutagenic mechanisms due to their remarkably similar profiles across all mutation types, but with EXO1 having a more significant role in the prevention of DSB-induced mutations.

**Figure 7:**
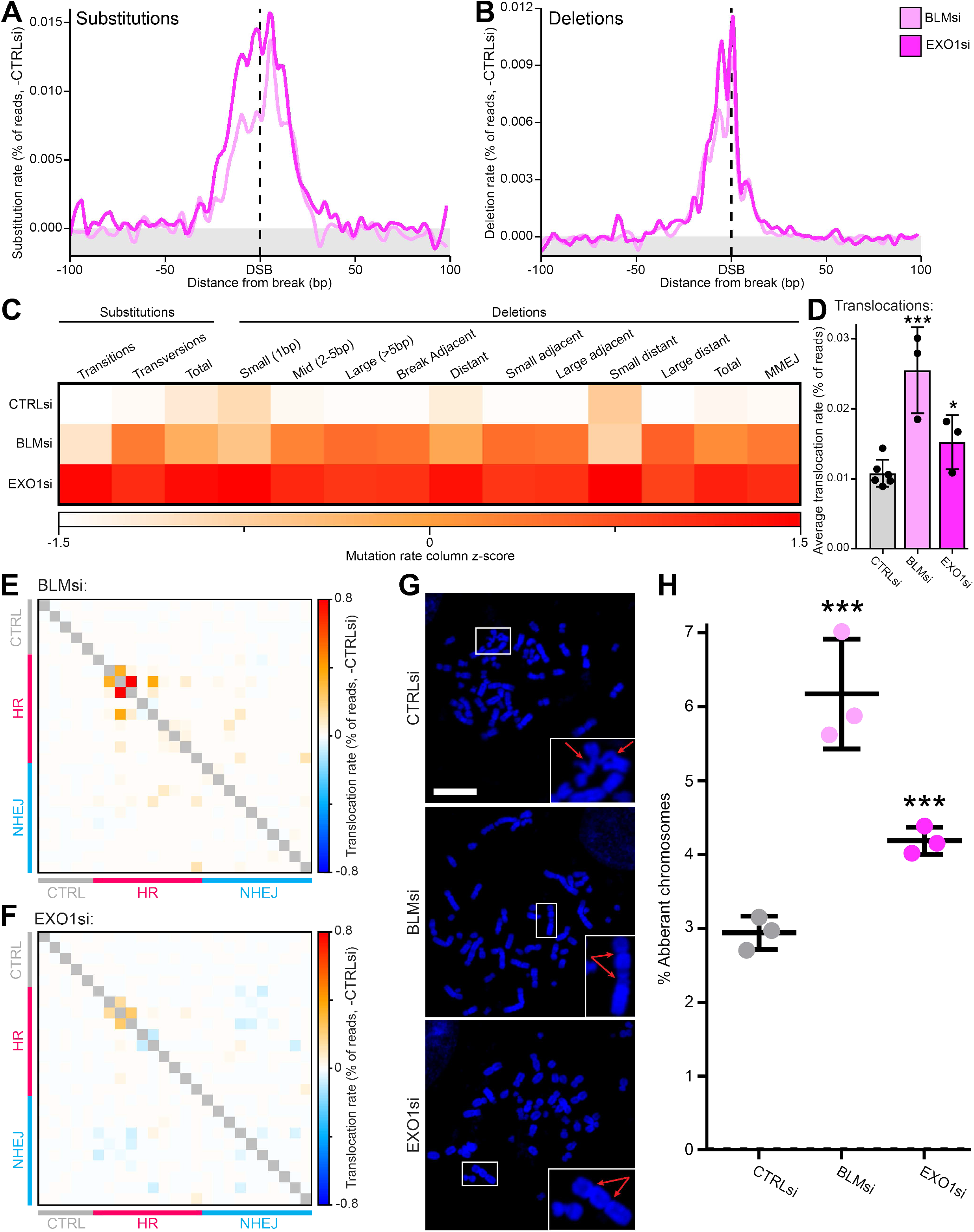
BLM and EXO1 synergistically protect against different mutations signatures. **(A)** Metagene line plot of base substitution rate as a percentage of readcount delta to -DSB then delta to control siRNA treatment, 100bp either side of AsiSI induced DSBs upon knockdown of BLM or EXO1, quantified by iMUT-seq. **(B)** Same as (A) but for deletion rate. (C) Heatmap of the average rate per DSB loci of different deletion types, calculated as a percentage of readcount delta to -DSB, quantified by iMUT-seq, and then converted into a column z-score to compare control, BLM and EXO1 siRNA treatment. **(D)** Average translocation rate between different DSBs quantified by iMUT-seq, with treatment using either control, BLM or EXO1 siRNA, points represent each biological replicate and error bars are S.D., all statistics done relative to CTRLsi using paired t-tests, * p<0.05, *** p<0.001. **(E)** Heatmap of translocation rates between different DSBs quantified by iMUT-seq with BLM depletion, each row and each column represent different iMUT-seq amplicons of either uncut control, HR-prone or NHEJ prone loci with each cell being the translocation rate as a percentage of total readcount delta to -DSB between the two loci that correspond to that cell and then delta to control siRNA. **(F)** Same as (E) but for EXO1 siRNA. **(G)** Representative images of metaphase spreads in HCT116 cells with 20μM etoposide treatment and with either control, BLM or EXO1 siRNA mediated depletion, scalebar is 10µm. Zoom in sections highlight chromosomal rearrangements, double arrows indicate dicentric chromosomes and single arrows indicate chromosome breaks. **(H)** Quantification of chromosomal aberrations in the metaphase spreads from (G) as a percentage of total chromosome number, points represent each biological replicate and error bars are S.D., statistics done relative to CTRLsi using a paired t-test, *** p<0.001.

However, translocation quantification showed that BLM knockdown increased translocations to a far greater degree than EXO1 knockdown (Figure 7D). BLM and EXO1 depletion exhibit slightly different translocation maps (Figure 7 E-F), although both primarily promote translocations between HR-prone loci (Figure S7B). Metaphase spread validation confirmed these results, clearly showing that whereas both EXO1 and BLM protect against chromosome aberrations, BLM has a far stronger role (Figure 7G-H). Interestingly, we noticed here that the rate of translocations mapped by iMUT-seq and the chromosomal rearrangements observed via metaphase spreads were very similar. A comparative analysis confirmed this, showing that control, BLM and EXO1 depletions all have similar relative changes in these two approaches (Figure S7C). This greatly supports iMUT-seq as an accurate approach in the mapping of translocations.

It has previously been hypothesised that different resection enzymes have varying functions at DSBs and cooperate to promote repair (Grabarz *et al*, 2013; Nimonkar *et al*, 2011; Sturzenegger *et al*, 2014; Tripathi *et al*, 2018), however it is still widely considered that these enzymes are mostly redundant (Sturzenegger *et al*, 2014; Gravel *et al*, 2008; Karanja *et al*, 2012). These results put this into questions and instead suggest that long-range resection enzyme choice has a direct link to the protection of the DNA against different mutations.

### Knockdown of Pol-δ and Pol-ε uncovers polymerase-error dependent DSB mutation signatures

As a further investigation of the mutations induced by disruption of resection, we depleted subunits of the major DNA polymerases used in HR repair; the Pol-δ subunit POLD1 and the Pol-ε subunit POLE to determine how these polymerases, that theoretically cause resection-dependent mutations via polymerase error, contribute to the mutational signature of DSBs.

Immediately we see that POLD1 depletion significantly increases distant base substitutions and deletions, while POLE depletion reduces these mutations to a very similar degree, almost mirroring the POLD1 results (Figure S8A-B). Analysis of the substitution signatures clearly shows that POLD1 knockdown increases C>T and T>C mutations, while decreasing T>A and T>G mutations, whereas POLE depletion only reduces C>T and T>C mutations (Figure S8C-D). This indicates that POLE is responsible for inducing the high levels of C>T and T>C mutations at DSBs, whereas POLD1 promotes lower levels of T>A and T>G errors. Both POLD1 and POLE depletion reduces large deletions at the break point, however POLD1 depletion also increases distant small deletions, whereas POLE depletion decreases these (Figure S8E-F) suggesting POLE is also responsible for slippage induced deletions at DSBs. Overall, it appears that Pol-δ is less prone to both base substitution errors and slippage-induced deletions than Pol-ε.

Surprisingly, translocation mapping finds that POLE depletion increases translocations whereas POLD1 depletion reduces them (Figure S8G-H). This indicates that, similar to our BLM and EXO1 depletion results, Pol-δ and Pol-ε protect against different mutation types, with Pol-δ maintaining sequence fidelity and Pol-ε protecting against chromosomal rearrangements.

### Differing roles of BRCA2, FANCA and RAD52 in preventing DSB-induced mutations

With the known role of BRCA2 and the potential role of FANCA in promoting RAD51 filament assembly, as well as the unclear role of RAD52 in mammalian DSB repair (Stark *et al*, 2004; Symington, Rothstein and Lisby, 2014; Kan, Batada and Hendrickson, 2017), a systematic analysis of the mutational signatures following depletion of these factors could provide significant insights into the mechanisms of late HR.

Depletion of RAD51 resulted in a precise spike of deletions on the break point as well as distant deletions and a broad base substitution peak (Figure 8A). Specifically, these deletions were predominantly small, mononucleotide deletions, though break adjacent large deletions also increased (Figure S9A). Knockdown of FANCA mimicked this profile, though at a significantly reduced level (Figure 8B, S9B), and knockdown of BRCA2 also resulted in increased distant deletions though surprisingly also resulted in a substantial increase in break adjacent large deletions (Figure 8C, S9C). RAD52 knockdown showed very little change in mutations, though did show a reduction in distant small deletions and a very subtle decrease in break adjacent large deletions (Figure 8D, S9D). Interestingly, no significant changes in base substitution signatures were observed with any of these knockdowns (Figure S9E).

**Figure 8:**
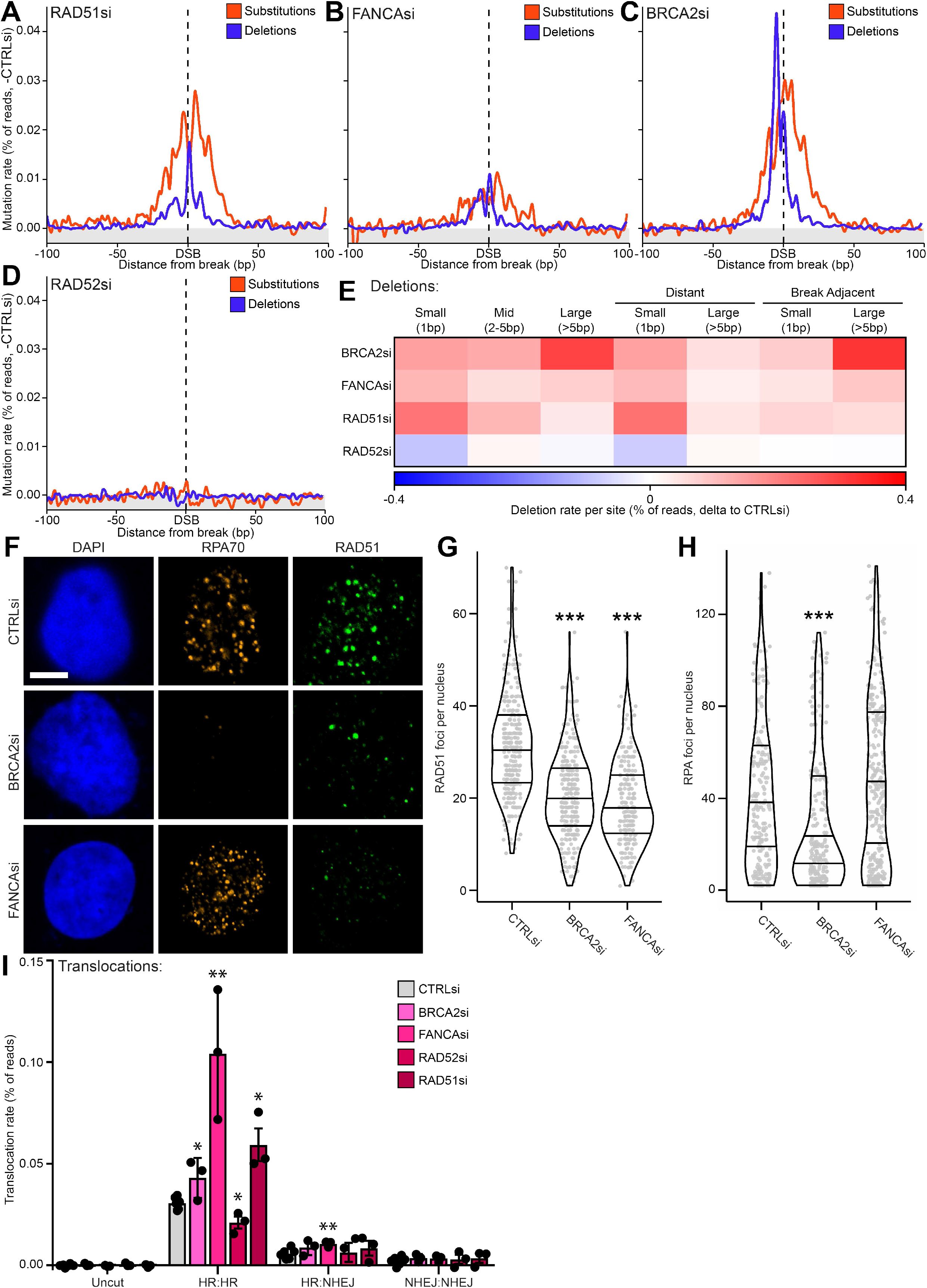
Differing roles of BRCA2, FANCA and RAD52 in preventing DSB-induced mutations. **(A)** Metagene line plot of base substitution and deletion rates as a percentage of readcount delta to -DSB and then delta of RAD51 siRNA – control siRNA, 100bp either side of AsiSI induced DSBs, quantified by iMUT-seq. **(B)** Same as (A) but for FANCA siRNA. **(C)** Same as (A) but for BRCA2 siRNA. **(D)** Same as (A) but for RAD52 siRNA. **(E)** Heatmap of the average rate per DSB loci of different deletion types, calculated as a percentage of readcount delta to -DSB then delta to control siRNA, quantified by iMUT-seq. **(F)** Immunofluorescence **of** U2OS cells treated with 5μM etoposide for 1hr followed by a 3hr recovery period, probing for RPA70 and RAD51 with either control, BRCA2 or FANCA siRNA treatment, scalebar is 5µm. **(G)** Quantification of RAD51 foci per nucleus in immunofluorescence from (F), lines represent median and interquartile ranges, statistics done using an unpaired Wilcoxon test, *** p <0.001. **(H)** Same as (G) but for RPA70 foci per nucleus. **(I)** Average translocation rate between different DSBs quantified by iMUT-seq, split by events between sites repaired by either HR or NHEJ or uncut control loci, with depletion of BRCA2, FANCA, RAD52 or RAD51, points represent each biological replicate and error bars are S.D., all statistics done relative to CTRLsi using paired t-tests, * p<0.05, ** p<0.01.

Both BRCA2 and FANCA depletions mimic the RAD51 depletion signature, however BRCA2 depletion has the additional signature of considerably increased large break-adjacent mutations (Figure 8E, S9C). This therefore suggests that BRCA2 and FANCA both contribute to RAD51 loading to different degrees, but that BRCA2 either deviates from this or has an additional role to this that leads to the increase in large deletions.

To investigate stage of the DDR, we conducted immunofluorescence in U2OS cells following etoposide treatment and stained for RAD51 alongside RPA70 as a marker of resected DNA to assess BRCA2 and FANCA contributions to RAD51 loading. Both BRCA2 and FANCA knockdown resulted in significantly reduced RAD51 foci per cell, indicating a significant role in promoting RAD51 recruitment (Figure 8F-G). Interestingly, whereas FANCA depletion had no effect on RPA70 focus formation, BRCA2 depletion significantly reduced RPA70 foci per cell (Figure 8F, H). This suggests that FANCA does not contribute to resection, but does promote RAD51 recruitment, whereas BRCA2 also promotes or maintains resection at DSBs, indicating an earlier role in DSB repair and could explain the additional large deletion phenotype we observed.

Surprisingly, translocation mapping tells a different story. Whereas both RAD51 and BRCA2 depletions cause a significant increase in translocations between HR-prone loci, FANCA knockdown results in a greater increase in translocations and also increased translocations at other loci (Figure 8I). This therefore indicates an alternative mechanism for FANCA in the prevention of translocations during DSB repair. Interestingly, RAD52 depletion causes a reduction in translocations between HR-prone loci (Figure 8I), which in combination with the reduction in small distant and large break-adjacent deletions supports the role of mammalian RAD52 in the mutagenic single-strand annealing (SSA) pathway (Stark *et al*, 2004; Kan, Batada and Hendrickson, 2017).

## DISCUSSION

Despite the significant role of DSB-induced mutagenesis in aging, cancer biology and other diseases, our understanding of these DSB-induced mutations is relatively limited, especially at the nucleotide level. Most studies that investigate mutations either use reporter assays, whole-genome sequencing or, in the investigation of genomic rearrangements, metaphase spreads. Whereas whole-genome sequencing can detect mutations at the nucleotide level across the genome, it is impossible to determine the location of the mutation relative to the initial damage, which is critical in determining the mutagenic mechanism, and although reporter assays can induce damage at known locations these are often exogenous sequences added to the genome (Ahrabi *et al*, 2016; Hussmann *et al*, 2021; Schep *et al*, 2021). In addition, both of these approaches normally require high levels of DNA damage (Kucab *et al*, 2019; Ahrabi *et al*, 2016; Hussmann *et al*, 2021; Schep *et al*, 2021) as they have relatively low sensitivity in determining mutation rates. To address this, we developed iMUT-seq, a technique that profiles mutations at extremely high sensitivity and at single nucleotide resolution around endogenous DSBs spread across the genome, allowing for the investigation of DSB-induced mutations at a level never seen before.

We initially compared the mutations at loci prone to either NHEJ or HR to elucidate repair pathway specific mutation signatures (Figure 2). Whereas both groups of loci showed similar mutations, likely due to most loci using both pathways under different cellular conditions, we found relative differences in the rates of different mutations between these loci. NHEJ-prone loci were relatively more susceptible to break-adjacent deletions (Figure 2D-F), whereas HR-prone loci were relatively more susceptible to base substitutions and mononucleotide deletions (Figure 2A-C, F). We therefore can infer that NHEJ leads to mutagenesis due to unrecovered degradation of the broken DNA ends, whereas HR induces mutations due to the need for re-polymerisation of the resected DNA, leading to polymerase error and slippage.

Our work here has profiled the mutational signature of 20 different DSB repair factor depletions, characterising the mutations induced by disruption of almost all aspects of the major DSB repair pathways. As well as provided remarkable insights into DSB-induced mutagenesis, we believe the results of our work here will act as a baseline to provide necessary context for future results using this technique. iMUT-seq can be used to examine novel repair factors and their mutational signatures can be mapped to the results here, allowing for a more advanced interpretation of the results. In addition, our findings have raised further questions that require additional investigation, such as the unexpected role of early NHEJ signalling in maintaining genome stability (Figure 4) or the mutational impact of mutagenic pathways such as MMEJ (Figure S2D-G, 5E).

Analysis of early NHEJ factors demonstrated the need for validation experiments, for which we recommend metaphase spreads to provide a global view of the genome as well as to validate translocation results (Figure 4E-I). Interference with certain repair processes, such as early NHEJ signalling, promotes massive destabilisation of the genome, leading to chromosomal loss which cannot be mapped by targeted sequencing experiments, but that are very apparent in metaphase spread analysis (Figure 4G-I).

The considerable increase in translocations upon LIG4 depletion (Figure 5F) was the greatest increase in translocations we observed. This is particularly interesting given the comparably small induction of translocations by XRCC4 depletion. XRCC4 depletion primarily promoted MMEJ repair, which is likely due to its role in promoting polymerase recruitment to fill in overhangs at DSBs (Craxton *et al*, 2018) which are required for MMEJ repair. This suggests that LIG4 may be recruited to XRCC4 once the DSB ends are ready for ligation and are therefore theoretically blunt. Thus, it is possible that in the absence of LIG4, DSBs are maintained that are prepared for ligation and are therefore easily ligated incorrectly to other DSB ends, leading to greatly increased translocations.

One of the more surprising results of these experiments was the mutagenic nature of HR repair. Although previously thought to prevent mutations, more recent studies have highlighted a mutagenic aspect of HR that is due to the need for repolymerisation of kilobases of DNA, leading to polymerase errors, and also the potential for misalignment of the damaged DNA on the sister chromatid which leads to expansions, contractions and even rearrangements (Guirouilh-Barbat *et al*, 2014; Hicks, Kim and Haber, 2010; Mosbach *et al*, 2020; Yang *et al*, 2008; Deem *et al*, 2011). This is greatly supported by our results here (Figure 6A-C, S6C-E), and we also demonstrated that HR has a significant role in preventing translocations (Figure 3D, 6D-G). Given the serious implications of translocations in mammalian biology, i.e. the strong association of chromosomal rearrangements in diseases such as cancer, it is possible that a primary function of HR is to reduce the rate of translocation at DSBs.

The differential mutational signatures of BLM and EXO1 depletion are of great importance to the understanding of long-range resection. Multiple previous studies have explored the roles of various resection enzymes (Karanja *et al*, 2012; Nimonkar *et al*, 2011; Sturzenegger *et al*, 2014; Tripathi *et al*, 2018), and although we limited our investigation to BLM and EXO1, the fact that we see such striking disparity in the induced mutation levels between the depletion of these two enzymes is critical (Figure 7). This result implies that different resection enzymes have overlapping, but not redundant roles, and instead function to prevent different mutagenic mechanisms from taking place. Some studies have suggested that different resection enzymes are more efficient in resecting DNA with nucleotide adducts or secondary structures (Grabarz *et al*, 2013; Nimonkar *et al*, 2011; Sturzenegger *et al*, 2014; Tripathi *et al*, 2018; Karanja *et al*, 2012), which could explain this result as these different obstructions could have various mutagenic impacts.

The mutational signature of BRCA2 depletion is a good example of how the analysis of these mutations can help elucidate complex mechanisms. Although BRCA2 displays a substitution, deletion and translocation signature of RAD51 (Figure 8C, I), the overlay of a large deletion signature suggests an alternative role (Figure 8E, S9C). This was further identified as a contribution to DNA resection since BRCA2 depletion reduced RAP70 foci. Previous reports have shown a role for BRCA2 in maintaining and promoting DNA end resection (Han *et al*, 2017; Chen, B. *et al*, 2021) and other studies have characterised a role for BRCA2 in a mechanism of RNA-dependent DSBR (D’Alessandro *et al*, 2018; Sessa *et al*, 2021; Bader and Bushell, 2020), and therefore this phenotype could be related to the mechanisms involving RNA at DSBs instead.

The extremely high sensitivity of iMUT-seq coupled to mapping of mutations at single-nucleotide resolution around DSBs has yielded novel insights into the mutations that arise around DSBs, the mutagenic mechanisms that derive them and also mechanistic details of the damage response as a whole. These novel insights into repair mechanics mark a significant step forward in our understanding of DSB repair and its mutagenic consequences. Additionally, significant new insight has been gained using iMUT-seq detailing the mechanisms involved in DSB-induced mutagenesis, which we believe has not only shown the utility of this technique, but we hope has inspired further research. Further study is needed to fully understand these mutagenic mechanisms; however the clear utility of this technology will be crucial for our further development towards understanding DSB-repair mechanisms. Finally, we believe our results are of considerable importance with regards to the development of DDR targeting chemotherapeutics, and future work should consider the potential off-target mutagenesis induced by these drugs in healthy cells, which although may not be acutely toxic, could lead to long-term complications or even future carcinogenesis. This technology could be applied to tailoring therapeutics towards limiting the incorporation of off-target mutations, reducing toxicity while maintaining efficacy.

## AUTHOR CONTRIBUTIONS

A.S.B. developed iMUT-seq, conducted all experiments and bioinformatic analysis and wrote the manuscript. A.S.B. and M.B. both conceived the technique and contributed to manuscript drafting.

## ACKNOWLEDGEMENTS

We thank Cancer Research UK for their core funding to the CRUK Beatson Institute A17196 and A31287 and for core funding for the Bushell lab (A29252). We also thank Gaëlle Legube and their lab for gifting us the AID-DIvA cells and for their continual support throughout the project.

## SUPPLEMENTAL FIGURE TITLES AND LEGENDS

**Figure S1: iMUT-seq technique and analysis design. (A)** Agilent Tapestation gel image of iMUT-seq genomic amplicons and iMUT-seq final library, the amplicons are an average of ∼270bp whereas the final library is ∼400bp due to the addition of the Illumina adapters. **(B)** Diagrammatic depiction of the machine learning approach used to optimise the iMUT-seq alignments. Several alignment parameters that alter how mismatches, deletions and insertions are handled by the aligner can be optimised to improve alignment efficiency. The machine learning tool started with several different combinations of the parameters to be optimised, and used a genetic algorithm to procedurally improve the alignment over multiple rounds of testing. After each round, the parameter combination with the highest alignment score was carried forward and a new set of combinations was generated based off of this high scoring combination. Therefore, after each round the alignment got increasingly effective, until the optimal parameter settings were achieved. **(C)** Comparison of alignment efficiency pre- and post-optimisation of the alignment using the machine learning approach from (B), each point represents an independent biological replicate of iMUT-seq and error bars are S.D. Pre-algorithm, the number of unaligned reads with DSB induction was significantly higher than without DSB induction, indicating that DSB-induced mutations are reducing alignment efficiency. Whereas post-algorithm, there was no significant difference in the number of unaligned reads between with and without DSB induction.

**Figure S2: iMUT-seq profiling of mutations at HR and NHEJ-prone DSBs. (A)** Metagene line plot of small and large deletion rate, as a percentage of readcount, 100bp either side of all AsiSI induced DSBs quantified by iMUT-seq **(B)** Same as (A) but for insertions at DSBs prone to repair by either NHEJ or HR, or at uncut control loci. **(C)** Boxplot of total deletions or insertions per loci as a percentage of readcount at either uncut control or DSB loci quantified by iMUT-seq, statistics done using a unpaired Wilcoxon test. **(D)** Same as (B) but for microhomology-mediated end-joining (MMEJ) rates, deletions are reported at the first nucleotide of the deletion resulting in the peak skewing to the left of the DSB. **(E)** Same as (C) but for MMEJ rates per loci at either uncut control, HR-prone or NHEJ-prone loci quantified by iMUT-seq. **(F)** Boxplot of deletion lengths at HR-prone or NHEJ-prone DSB loci for either total deletions or specifically deletions that show microhomologies. **(G)** Boxplot of microhomology lengths at DSB loci prone to either NHEJ or HR repair. **(H)** Same as (E) but for translocation rates, statistics done via unpaired Wilcoxon test, * p<0.05. **(I)** The induction rate of mutations at DSBs, quantified by iMUT-seq, compared to their statistical significance, statistics done by paired t-tests comparing the mutations at each nucleotide around the break at DSB loci compared to uncut control loci, grey box indicates the significance threshold equivalent to a p-value of 0.05, zoom in section on the right with a dotted line at the estimated sensitivity limit of iMUT-seq of 0.0005% or 1 in 200,000. **(J)** Comparison of the rate of three deletion events from (I) at NHEJ-prone, HR-prone and uncut control loci with a dotted line at the estimated sensitivity limit of iMUT-seq of 1 in 200,000, points represent each biological replicate and error bars are S.D., statistics done relative to uncut by paired t-tests, * p<0.05, ** p<0.01, *** p<0.001.

**Figure S3: Interrupting DSB repair causes increasing mutation rates the later the interruption occurs in the pathway. (A)** Western blot validation of the knockdown of 19 different DSB repair factors from both NHEJ and HR repair, in AID-DIvA cells with and without OHT treatment for 4 hours to induce DSBs. Due to the large number of samples, each blot had to be spread across 3-5 gels, and to maintain comparability all blots for the same target were ran, probed and scanned together, then treated identically during image processing, and lines were added to define the separate gels. Phosphorylated ATM was probed as a measure of DSB induction and one of three loading controls; GAPDH, Lamin-A/C or Vinculin was used during each western blot. **(B)** Boxplot of iMUT-seq deletion lengths upon knockdown of 19 different DSB repair factors, all statistics are done relative to the control siRNA result using an unpaired Wilcoxon test, * p<0.05, ** p<0.01, *** p<0.001.

**Figure S4: DNA-PK inhibition causes a substantial increase in large deletions at DSBs. (A)** Metagene line plot of base substitution and deletion rates as a percentage of readcount delta to -DSB and then delta of DNA-PKi (10μM NU7441) – mock treatment, 100bp either side of AsiSI induced DSBs, quantified by iMUT-seq. **(B)** Bar plot of the rate of different deletion lengths per DSB loci as a percentage of readcount, quantified by iMUT-seq, with or without DNA-PKi treatment, points represent each biological replicate and error bars are S.D., statistics done using a paired t-test, * p<0.05, *** p<0.001. **(C)** Heatmap of translocation rates between different DSBs quantified by iMUT-seq with DNA-PKi treatment, each row and column represent a different iMUT-seq amplicon, each cell representing the translocation rate calculated between the row/column amplicons as a percentage of readcount delta to -DSB and then delta to mock treatment.

**Figure S5: Disruption of late NHEJ processes promotes large deletions and global translocations. (A)** Stacked bar plot of the relative base substitution rates per DSB loci quantified by iMUT-seq upon depletion of either POLL, XRCC4 or LIG4. **(B)** Bar plots of base substitution rates at either uncut loci or DSB loci prone to either NHEJ or HR repair, comparing CTRLsi to POLLsi, left shows substitutions adjacent to the break point and right shows substitutions distant from the break site, points represent each biological replicate and error bars are S.D., statistics done using a paired t-test, * p<0.05, ** p<0.01. **(C)** Bar plot of the rate of different deletion lengths per DSB loci as a percentage of readcount, quantified by iMUT-seq, with siRNA depletion of either POLL, XRCC4 or LIG4, points represent each biological replicate and error bars are S.D., statistics done relative to CTRLsi or between XRCC4si and LIG4si using a paired t-test, ** p<0.01, *** p<0.001. **(D)** Average translocation rate between different DSBs quantified by iMUT-seq, split by events between sites repaired by either HR or NHEJ or uncut control loci, treated with either control, POLL, XRCC4 or LIG4 siRNA, points represent each biological replicate and error bars are S.D., all statistics done relative to CTRLsi using paired t-tests, ** p<0.01, *** p<0.001.

**Figure S6: Early HR processes promote deletions at DSBs. (A)** Heatmap of the average rate per DSB loci of different mutation types after treatment with either ATM inhibition (10μM KU55933), ATR inhibition (10μM, VE-821) or combined ATM/ATR inhibition (10μM KU55933 + 10μM VE-821), calculated as a percentage of readcount delta to -DSB then delta to mock treatment, quantified by iMUT-seq. **(B)** Stacked bar plot of the relative base substitution rates per DSB loci quantified by iMUT-seq upon depletion or inhibition of either MRE11, ATM/ATR or BRCA1. **(C)** Metagene line plot of small (1bp) and large (>5bp) deletion rates as a percentage of readcount delta to -DSB and then delta of MRE11 siRNA – control siRNA, 100bp either side of AsiSI induced DSBs, quantified by iMUT-seq. **(D)** Same as (C) but for ATM/ATR inhibition. (E) same as (C) but for BRCA1 depletion.

**Figure S7: BLM preferentially protects against translocations compared to EXO1. (A)** Stacked bar plot of the relative base substitution rates per DSB loci quantified by iMUT-seq upon depletion of either BLM or EXO1. **(B)** Average translocation rate between different DSBs quantified by iMUT-seq, split by events between sites repaired by either HR or NHEJ or uncut control loci, with depletion of BLM or EXO1, points represent each biological replicate and error bars are S.D., statistics done relative to CTRLsi using paired t-tests, * p<0.05, *** p<0.001. **(C)** Comparison of the rate of DSB-induced chromosomal rearrangements quantified by either metaphase spread or iMUT-seq in cells treated with either control, BLM or EXO1 siRNA, points represent each biological replicate and error bars are S.D.

**Figure S8: Knockdown of Pol-δ and Pol-ε defines polymerase error-dependent DSB mutation signatures. (A)** Metagene line plot of base substitution rate as a percentage of readcount delta to -DSB then delta to control siRNA, 100bp either side of AsiSI induced DSBs upon knockdown of POLD1 or POLE, quantified by iMUT-seq. **(B)** Same as (A) but for deletions. **(C)** Heatmap of the average rate of each base substitution type per DSB loci quantified by iMUT-seq, with treatment using either control, POLD1 or POLE siRNA. **(D)** Stacked bar plot of the relative base substitution rates per DSB loci quantified by iMUT-seq upon depletion of either POLD1 or POLE. **(E)** Metagene line plot of small (1bp) and large (>5bp) deletion rates as a percentage of readcount delta to -DSB and then delta of POLD1 siRNA – control siRNA, 100bp either side of AsiSI induced DSBs, quantified by iMUT-seq. **(F)** Same as (E) but for POLE siRNA. **(G)** Heatmap of translocation rates between different DSBs quantified by iMUT-seq with POLD1 depletion, each row and column represents a different iMUT-seq amplicon, each cell representing the translocation rate calculated between the row/column amplicons between the row/column amplicons as a percentage of readcount delta to -DSB and then delta to control siRNA. **(H)** Same as (G) but for POLE siRNA.

**Figure S9: The effects of depletion of BRCA2, FANCA, RAD52 and RAD51 on DSB-induced deletion lengths and base substitution signatures. (A)** Metagene line plot of small (1bp) and large (>5bp) deletion rates as a percentage of readcount delta to -DSB and then delta of RAD51 siRNA – control siRNA, 100bp either side of AsiSI induced DSBs, quantified by iMUT-seq. **(B)** Same as (A) but for FANCA siRNA. **(C)** Same as (A) but for BRCA2 siRNA. **(D)** Same as (A) but for RAD52 siRNA. **(E)** Stacked bar plot of the relative base substitution rates per DSB loci quantified by iMUT-seq upon depletion of either BRCA2, FANCA, RAD52 or RAD51.

## STAR METHODS

### Resource availability

#### Lead contact

Further information and requests for resources and reagents should be directed to and will be fulfilled by the lead contact, Martin Bushell

#### Materials availability

This study did not generate new unique reagents.

#### Data and code availability

All iMUT-seq raw data has been deposited at ArrayExpress and are publicly available at the date of publication.

All analytical code is publicly available on GitHub (https://github.com/aldob/iMUT-seq), which includes the raw data processing pipeline and it’s parameters as well as the code used to generate plots in R.

Any additional information required to reanalyse the data reported in this paper is available form the lead contact upon request.

### Experimental model and subject details

#### Cell culture

U2OS, AID-DIvA and HCT116 cells were all cultured in Dulbecco Modified Eagle’s Medium (DMEM, GibCo) supplemented with 10% Fetal bovine serum and 2mM L-glutamine, with AID-DIvA cell culture medium also supplemented with 800 µg/mL G418 (Formedium, G418S). All cells were incubated at 37°C with a 5% CO2, humidified atmosphere. U2OS and AID-DIvA cells are female and HCT116 cells are male. All cell lines were tested for mycoplasma contamination each time they entered cell culture from storage and were always found to be negative.

### Method details

#### Cell transfection

All transfections were completed using Dharmafect 1 (Horizon Discovery, T-2001-03). Dharmafect 1 was used at a final dilution of 1/1000 and siRNA (Table S2) was used at a final concentration of 20nM. Both Dharmafect and siRNA were separately diluted in serum free medium to a volume that was 5% of the desired final medium volume and incubated at RT for 5mins. The siRNA and Dharmafect were then combined in a 1:1 ratio and incubated for 20mins at RT. The siRNA dilution was then made up to the desired final volume with antibiotic free medium, and the cell culture medium was immediately replaced with this transfection medium. Cells were then incubated for 24 hours, after which the medium was refreshed with regular, antibiotic free medium. Cells were then incubated for a further 48 hours before experimental treatments began.

#### Cell treatments

For DSB induction in AID-DIvA cells, treatment with 300nM 4-hydroxytamoxifen (OHT) was given for 4 hours. For DSB repair via degradation of the AsiSI fusion protein, OHT treated cells were washed twice in pre-warmed PBS, then once in pre-warmed medium containing 500 µg/mL auxin (IAA) (Sigma, I5148) and then finally replaced with fresh medium containing 500 µg/mL auxin and incubated for 14 hours.

ATMi (KU-55933, Merck SML1109), ATRi (VE-821, Merck SML1415) and DNA-PKi (NU7441, Tocris 3712) were all used at 10μM and administered in a 1 hour pre-treatment and maintained throughout the period of the experiment.

Etoposide treatments were completed for 1 hour with either 5µM or 20µM concentration. Cell culture medium was replaced with pre-warmed medium containing the desired concentration of etoposide, incubated for 1 hour and then cells were washed twice in pre-warmed PBS, once in pre-warmed medium and finally replaced with fresh medium and incubated for the indicated recovery period.

#### Western Blotting

Cells were lysed in 1.1x NuPAGE LDS sample buffer (Thermofisher, NP0007), scraped from their plates and transferred to microcentrifuge tubes. Samples were passed through a 23 gauge needle 10 times to shear DNA and homogenise the samples, and then heated to 95C for 10 minutes. Samples were run on either 4-12% NuPage Bis-Tris gels (Thermofisher, NP0322PK2) or 10% polyacrylamide gels. And transferred in tris-glycine transfer buffer containing 20% methanol and 0.05% SDS onto nitrocellulose membranes for 1.5 hours at 100V. Membranes were then blocked in TBST containing 5% BSA at RT for 1 hour. Primary antibody probing was done overnight at 4C with all antibodies diluted in TBST containing 5% BSA. Membranes were then washed three times in TBST for 10 minutes at RT and then probed with secondary antibodies (Li-COR Biosciences) diluted 1/10000 in 5% BSA in TBST for 1 hour at RT. Membranes were then washed three times in TBST for 10 minutes at RT before scanning with a Li-COR Odyssey.

#### Metaphase spreads

500,000 HCT116 cells were seeded onto 10cm plates and incubated for 24 hours before being transfected as described above. Transfected cells were incubated for 24 hours, then their medium was replaced, and they were incubated for a further 48 hours. The cells were then treated with 20μM etoposide for 1 hour, washed twice with pre-warmed PBS and once with pre-warmed medium and then allowed to recover for 14 hours. Metaphase cells were then enriched by treating with the microtubule poison colcemid (Sigma, D7385) at 200nM for 1 hour.

Cells were trypsinised, pelleted, washed once in PBS and then re-suspended in 10mL of 75mM potassium chloride. Cells were then allowed to swell by incubating at 37°C for 30 minutes. 5mL of ice-cold fixative (75% methanol, 25% acetic acid) was then slowly added to the cells. Cells were then pelleted at 200g, resuspended in 10mL of fixative and incubated on ice for 2 mins twice to completely fix and wash off any buffer from the cells. Cells were then finally pelleted and resuspended in 5mL of fixative and dropped onto glass slides from a height of 15-20cm using a p200 pipette. Slides were then steamed for 5s over a water bath set to 80°C, and then checked under a light microscope to ensure optimal spreading before drying overnight. The slides were then stained with DAPI by immersing in water containing 0.1μg/mL DAPI, washed by immersing in water and then allowed to dry overnight. Coverslips were then mounted to the slides with Vectashield anti-fade mounting medium (Vector Laboratories, H-1000). Spreads were then imaged using a Carl-Zeiss LSM 710 confocal microscope under a 63x objective and analysed in ImageJ.

#### Immunofluorescence

4000 U2OS cells were seeded onto 12-well removable silicon chambered slides from Ibidi (Thistle Scientific, 81201) and transfected as described earlier. Cells were then treated with 5μM etoposide for 1 hour before washing twice with pre-warmed PBS and once with pre-warmed medium and then allowed to recover for 3 hours. Cells were then pre-extracted at RT for 3min in CSK buffer (100mM NaCl, 300mM sucrose, 3mM MgCl2, 10mM PIPES pH 7.0, 50mM NaF, 5mM sodium orthovanadate, 10mM β-glycerol phosphate and 0.7% Triton). Cells were washed once in CSK, once in PBS and fixed in PBS containing 2% paraformaldehyde for 10 minutes at RT. Cells were washed once in PBS, once in TBST and blocked with TBST containing 10% goat serum (Merck, G9023) for 1 hour at RT. Cells were washed twice in TBST and probed overnight at 4°C with primary antibodies diluted in TBST containing 1% goat serum. Cells were washed 4 times in TBST for 5mins at RT and probed with alexa-fluor conjugated secondary antibodies, diluted 1/1000 in TBST with 1% goat serum, for 1 hour at RT. Slides were washed 4 times in TBST for 5mins at RT, dipped in water to remove residual buffer, and a coverslip was mounted using Vectashield anti-fade hard-set mounting medium containing DAPI (Vector Laboratories, H-1500). Slides were imaged using a Carl Zeiss LSM 710 confocal microscope under a 63x objective and images were analysed in Fiji(Schindelin *et al*, 2012) using the FindFoci plugin(Herbert, Carr and Hoffmann, 2014).

#### iMUT-seq experimental protocol

All iMUT-seq experiments were done in three biological replicates that were independently carried out. 60,000 AID-DIvA cells were seeded into 6-well plates, incubated for 24 hours and transfected as described earlier with siRNA from Table S2. Transfected cells were incubated for 24 hours, their medium was replaced, and they were incubated for a further 48 hours. Cells were treated with or without 300nM 4-hydroxytamoxifen (OHT) for 4 hours to induce DSBs, washed twice with pre-warmed PBS and once with pre-warmed medium containing 500μg/mL IAA, and replaced with media containing 500μg/mL IAA and incubated for 14 hours to degrade the AsiSI fusion protein and allow complete DSB repair. Cells were placed on ice, washed once in PBS and lysed in cytoplasmic lysis buffer (50mM HEPES pH7.9, 10mM KCl2, 1.5mM MgCl2, 0.34M sucrose, 0.5% triton, 10% glycerol, 1mM DTT) for 10 minutes. Cells were washed once in cytoplasmic lysis buffer and the nuclei were lysed in genomic extraction buffer (50mM Tris pH 8.0, 5mM EDTA, 1% SDS, 0.5mg/mL Proteinase K). The nuclear lysates were transferred to 2mL DNA LoBind microcentrifuge tubes (Fisher Scientific, 0030108426) and incubated in a thermomixer at 60°C for 40 minutes with 500rpm agitation. 0.1 volumes of 3M sodium acetate pH 5.2 was added followed by 2.5 volumes of 100% ethanol, the tubes were inverted several times to mix an then incubated on ice for 1 hour to precipitate the genomic DNA. The DNA was pelleted at 19000g for 20 minutes at 4°C, washed in 75% ethanol and re-pelleted for 10 minutes twice. The ethanol was aspirated and the DNA pellet allowed to dry at RT before resuspended in water and quantifying the DNA concentration via nanodrop.

For each condition, 3 50μL PCR reactions were run each containing 0.7μL Phusion polymerase (NEB, M0530L), 1X Phusion HF buffer, 2M betaine, 1.5% DMSO, 400μM dNTPs, multiplexed 25 genomic primer pairs (Table S2) at a concentration of 40nM per primer, i.e. 2µM total, 1μg genomic DNA. The reaction was carried out as follows: 98°C 5mins, 12 cycles of 98°C 60s, 62°C 120s, 72°C 120s, then 72°C 5mins. All 3 reactions were combined and 100μL of the total reaction volume was taken forward for size selection. 60μL of SPRISelect beads (Beckman, B23318) was added to the 100μL PCR reaction mixture, mixed by pipetting and then incubated for 5mins at RT. The beads were collected on a magnetic rack and the supernatant was transferred to a fresh tube. 70μL of beads were added to the supernatant and mixed by pipetting and incubated for 5mins at RT, binding the amplicons. Beads were collected on the magnet, the supernatant was discarded and the beads were washed twice in 85% ethanol for 30s. The beads were dried until most ethanol had evaporated, but the beads were still wet, and the beads were then resuspended in 50μL 10mM Tris pH 8.0 and incubated for 10 minutes at 37C with 1000rpm agitation. The 50µL of eluted amplicons were then transferred to a fresh tube and the process of bead selection was repeated but with half the volumes (30μL and 35μL of beads) and eluted in 25μL of 10mM Tris pH 8.0. 10μL of these amplicons were used for final library preparation using the NEB Ultra II DNA library prep kit according to the manufacturer’s instructions using a 1/25 adapter dilution and 5 PCR cycles.

These final libraries were then quantified using the Qubit dsDNA HS kit (Thermo, Q32854), pooled and sequenced on a NextSeq 500 using a high output 300 cycle kit set to run paired-end 150 cycles.

### Quantification and statistical analysis

#### iMUT-seq translocation mapping

Translocation mapping was conducted on raw fastq files using our custom tool mProfile TransloCapture (Available at https://github.com/aldob/mProfile and for install via https://pypi.org/project/mProfile-mut/). TransloCapture uses the sequences of the genomic primers initially used to amplify our target sequences to identify which primers were used to amplify each sequencing read, determining if the read is from an accurately repaired or a translocated DSB and which sites were translocated together. TransloCapture also allows the non-translocated and the translocated reads to be separate from the other reads and both to be written separately to new fastq files. TransloCapture outputs translocation map tables which were then used for all downstream translocation analysis via custom python and R scripts (See Key Resources table).

#### iMUT-seq mutation profiling

Exact parameters and settings used for raw data processing can be found in the raw data pipeline shell script (See Key Resources table). First, TransloCapture was used to filter out any translocated reads from the fastq files and Fastp(Chen, S. *et al*, 2018) was used to filter out low-quality reads prior to alignment with Bowtie 2(Langmead and Salzberg, 2012). Bowtie 2 parameters were determined via machine learning with the custom alignment machine python script available on GitHub, which yielded the following parameters: “--fr -- maxins 400 --no-discordant --no-mixed --ignore-quals --no-1mm-upfront -D 100 -R 50 -L 28 - N 1 --np 0 --dpad 49 --gbar 2 --mp 3.2,0.35 --rdg 1,1 --rfg 5,2 --score-min L,-1.0,-0.5”. Alignments were then sorted using Samtools(Li, H. *et al*, 2009) sort and indexed using Samtools index. Raw mutation calls were then generated using Samtools mpileup. These raw mutation calls were then parsed by our custom tool mProfile callMUT (Available at https://github.com/aldob/mProfile and for install via https://pypi.org/project/mProfile-mut/) into mprofile mutation call files.

Mprofiles were then used for all downstream mutation analysis and quantification via custom python and R scripts (See Key Resources table).

#### iMUT-seq mutation quantification

All mutations are calculated as a percentage of reads at the nucleotide position that the mutation occurs. A delta of damaged-undamaged mutation rates was used to remove background mutations that are either naturally present within the genomes of our cells or that were induced via PCR or sequencing error. This delta was conducted on a per-nucleotide level, subtracting the rate of each mutation type at each individual genomic position in the undamaged sample from the rate of that mutation type at the corresponding position in the damaged sample. This generated a DSB-induced mutation profile for each condition. Where results are shown relative to control siRNA treatment, this same approach was taken to subtract the mutation rates for the damaged-undamaged delta of the control siRNA from the damaged-undamaged delta of the treatment siRNA at each nucleotide sequenced.

The mutation profiles, in the form of mprofile files, were then used either for directly generating metagene line plots of the mutation profiles, or to create average overall mutation rates. Where average mutation rates are used, this is calculated per site i.e. the average total mutation rate across the regions sequenced per DSB quantified.

#### Statistical analysis

All statistical tests used are detailed in the figure legends where they have been used. In all cases *p<0.05, **p<0.01, ***p<0.001. All experiments were conducted in biological triplicate, in which each was completed independently of the others, and where appropriate pairing of these replicates was used in statistical tests.

## SUPPLEMENTAL INFORMATION

Table S1. PCR primer sequences used for generation of the iMUT-seq libraries and associated metadata such as amplicon length. Related to Figures 1-8 and S1-9.

Table S2: siRNA sequences used for the screen of DSB repair factors and their sources. Related to Figures 3-8 and S3-9.

