## Supplementary figures and images for "iMUT-seq: high-resolution mapping of DSB mutational landscapes reveals new insights into the mutagenic mechanisms of DSB repair"

### Supplementary Figures

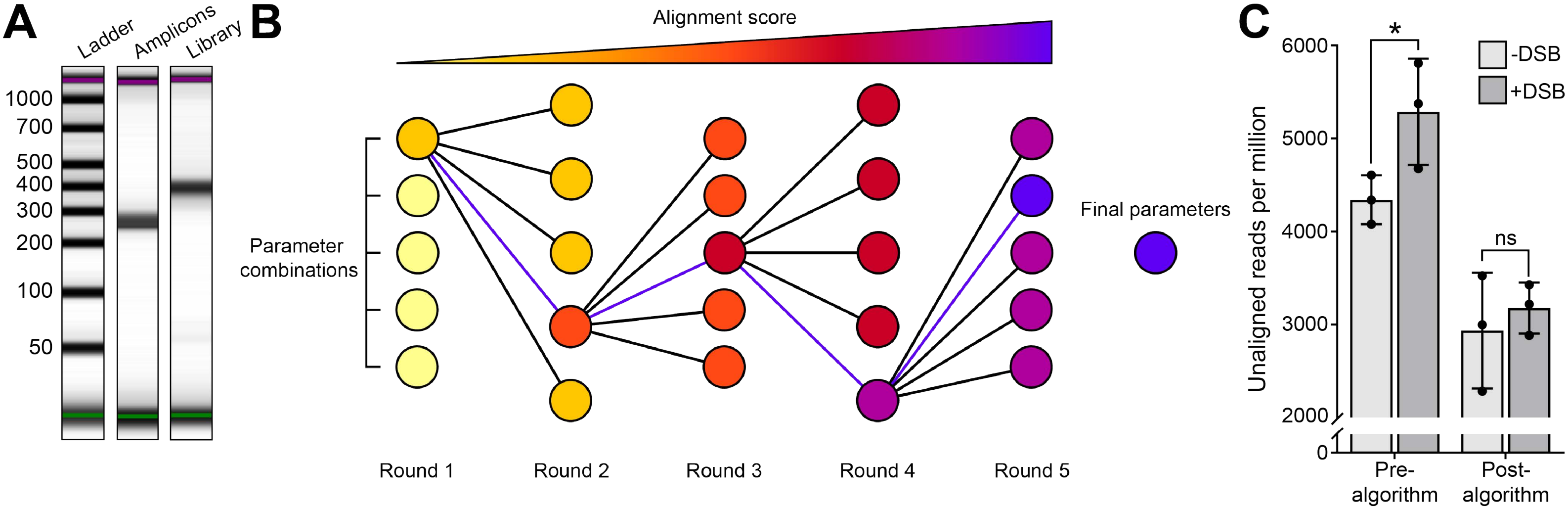

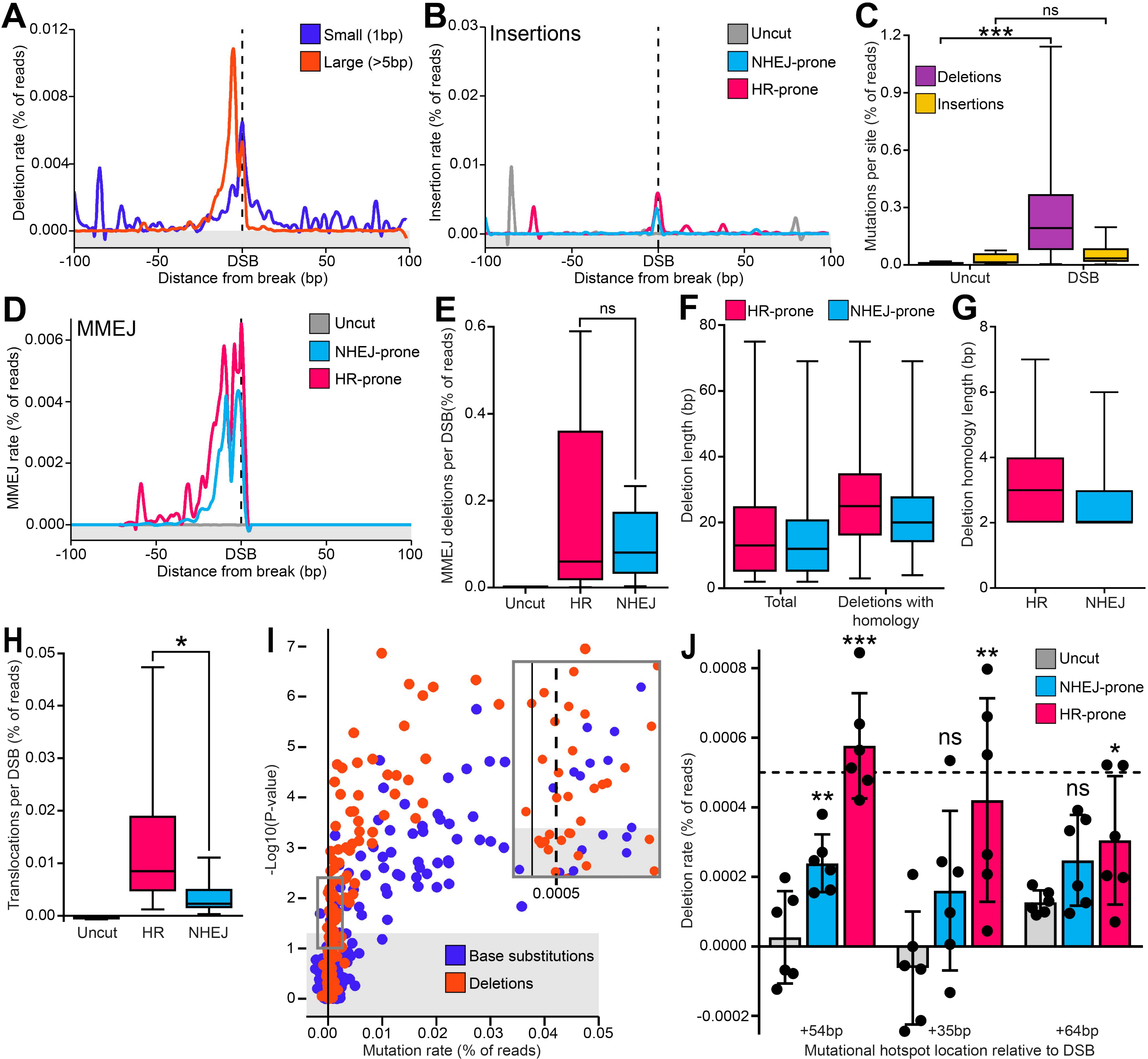

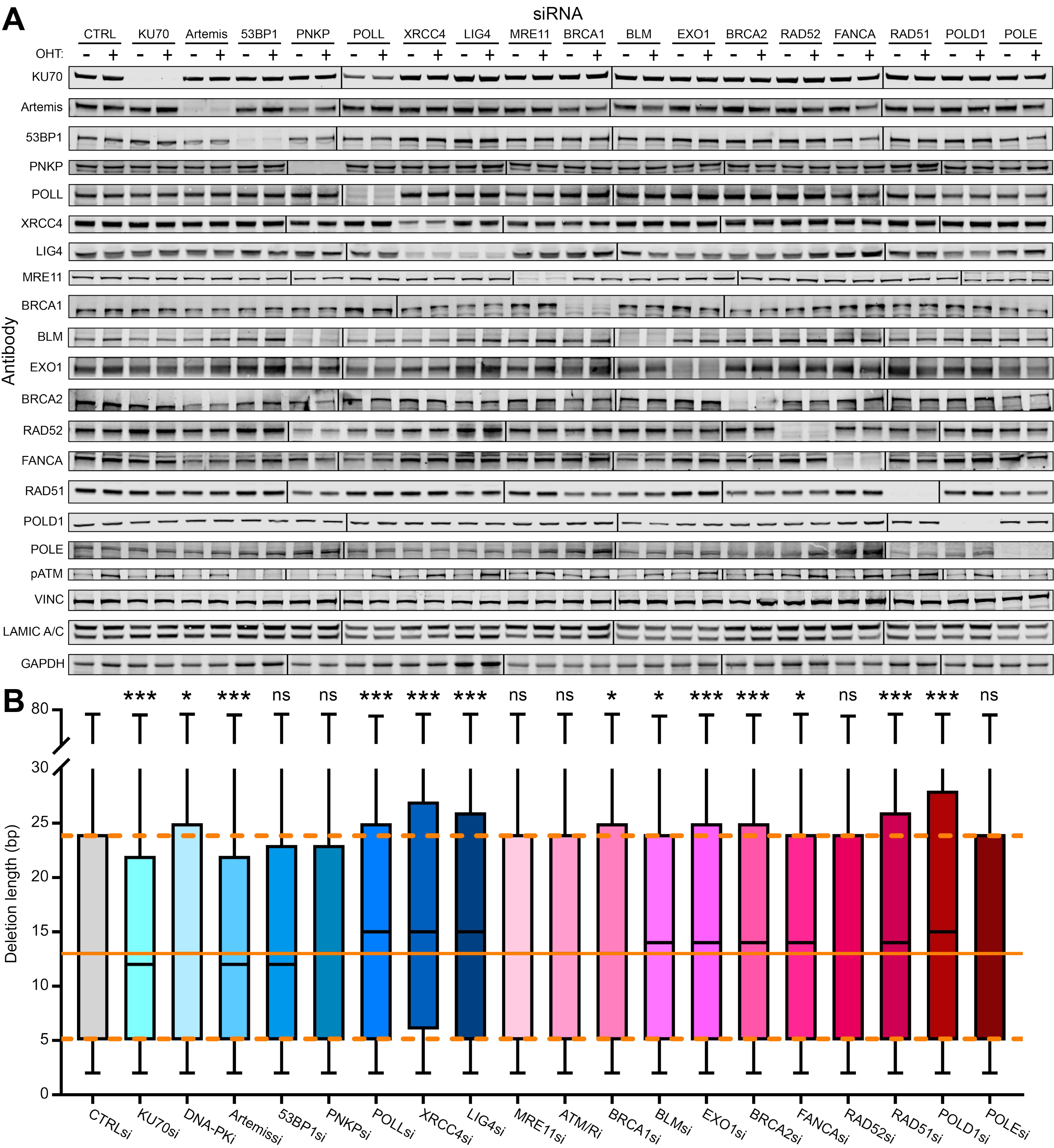

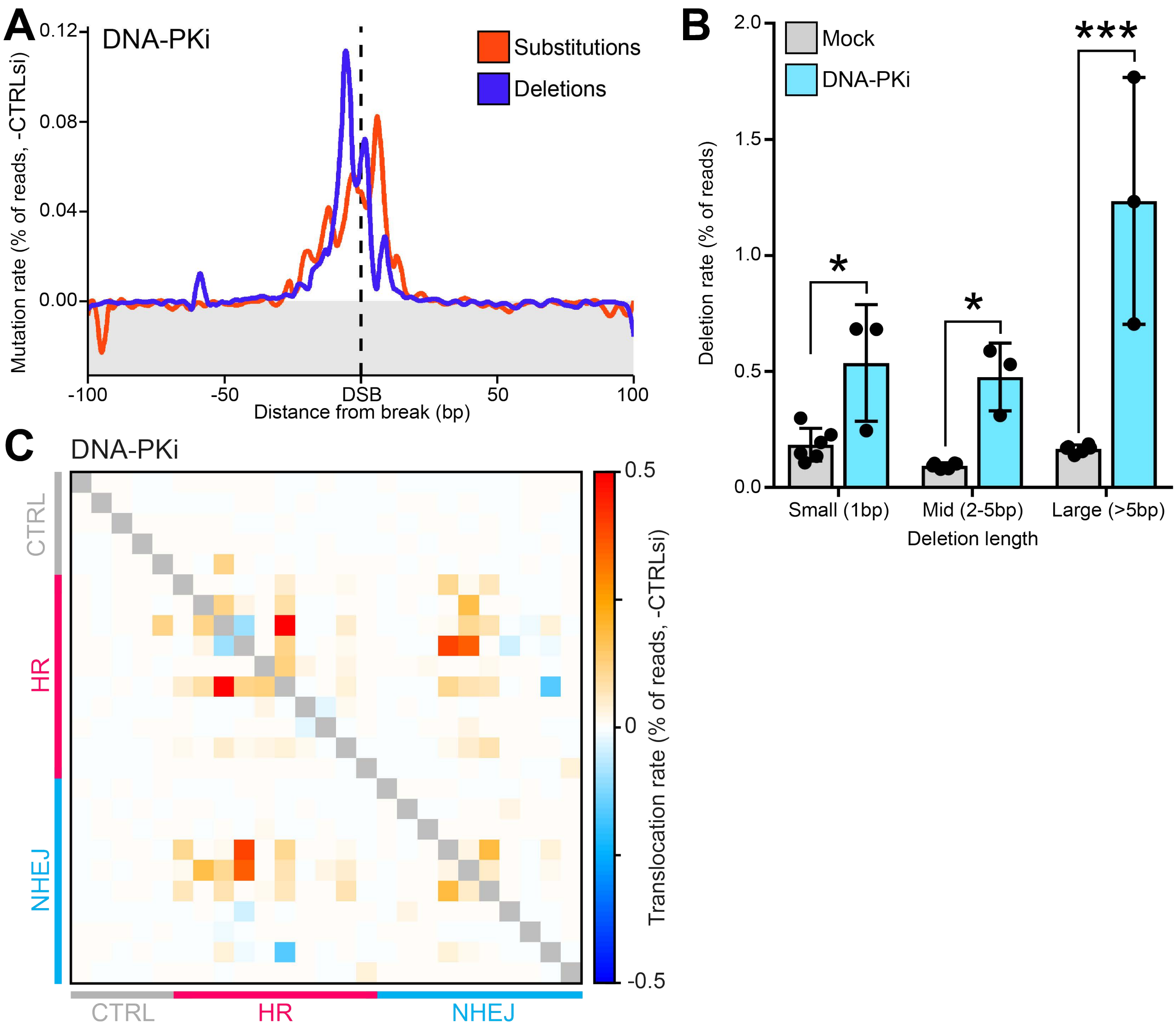

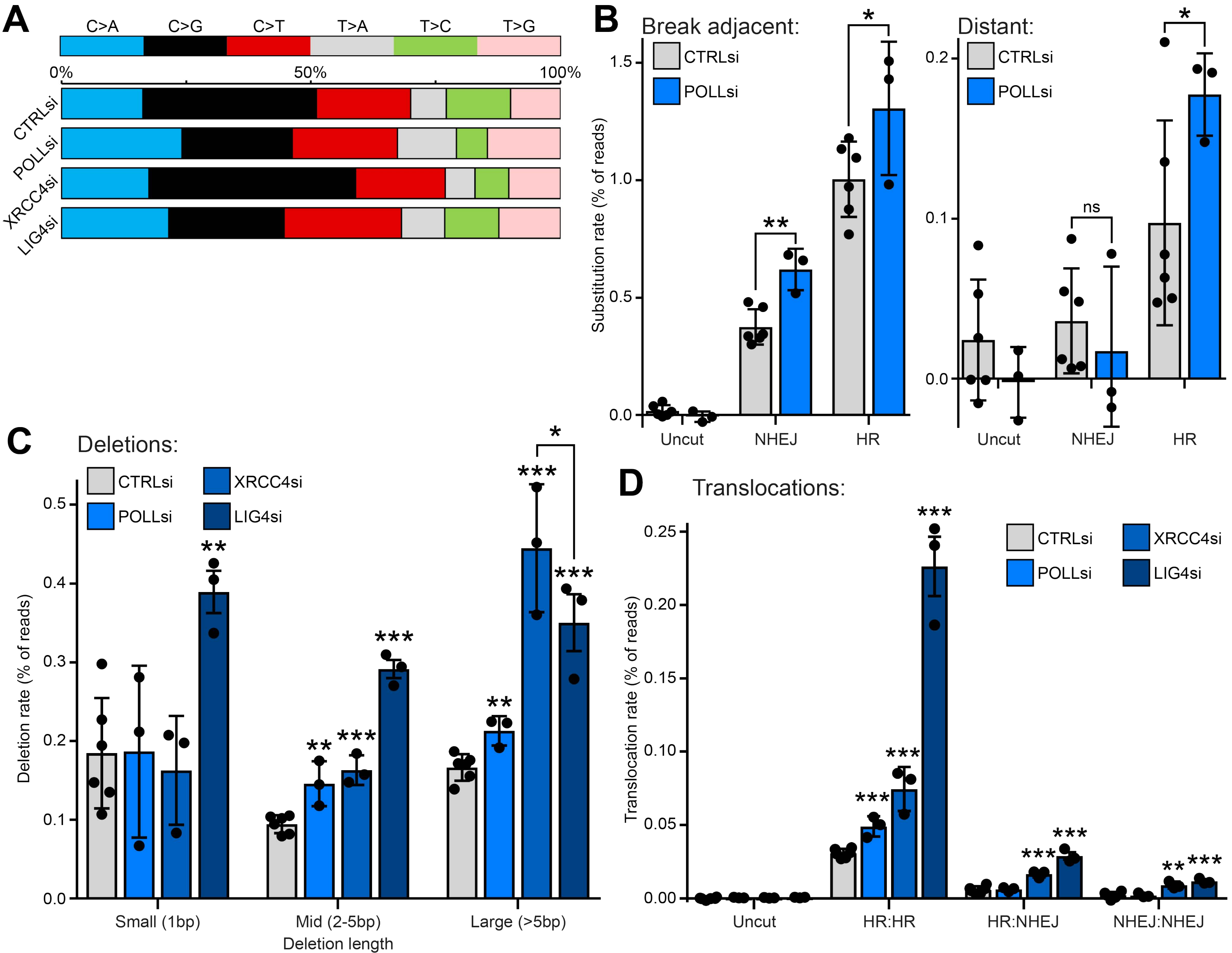

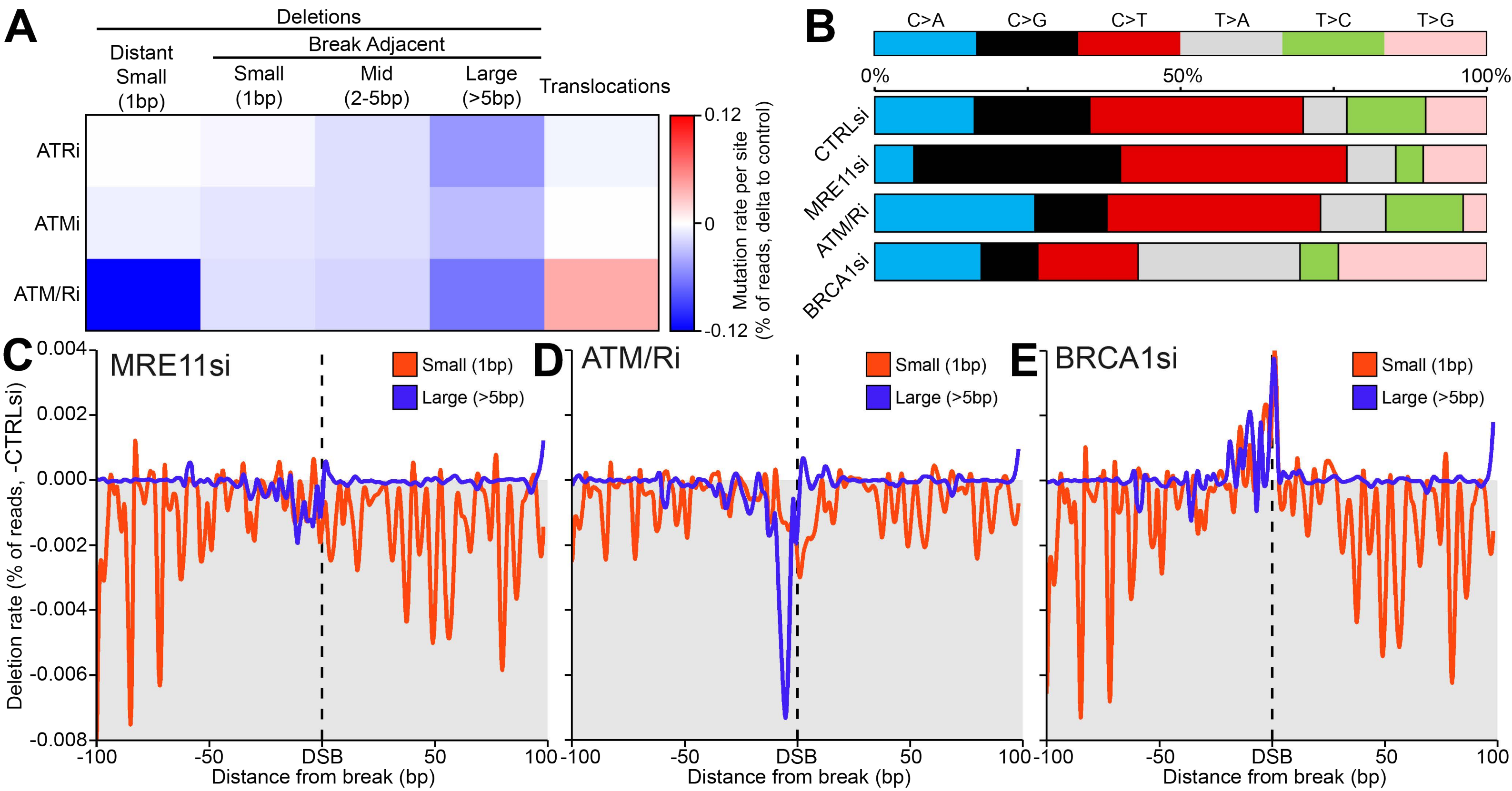

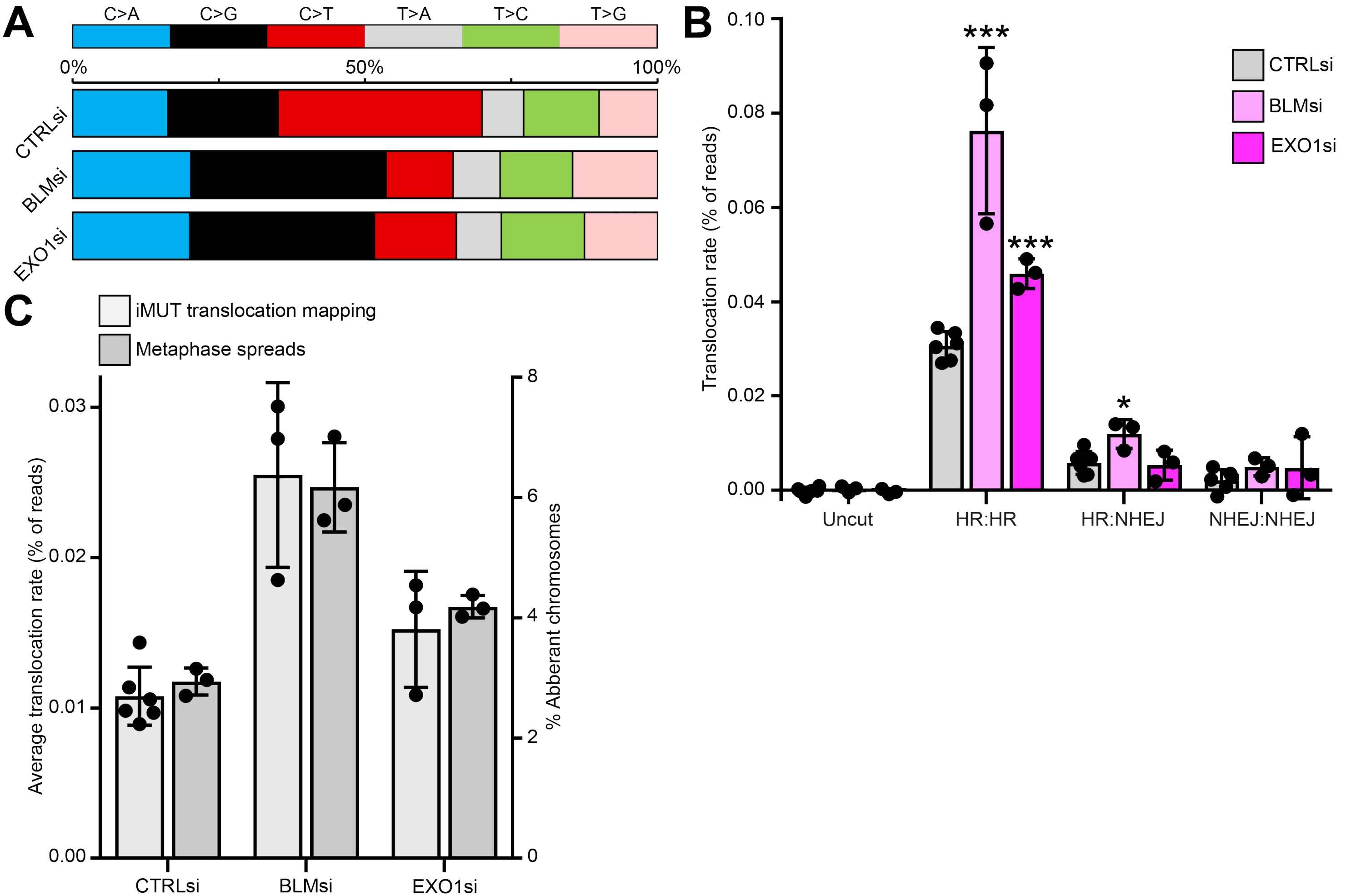

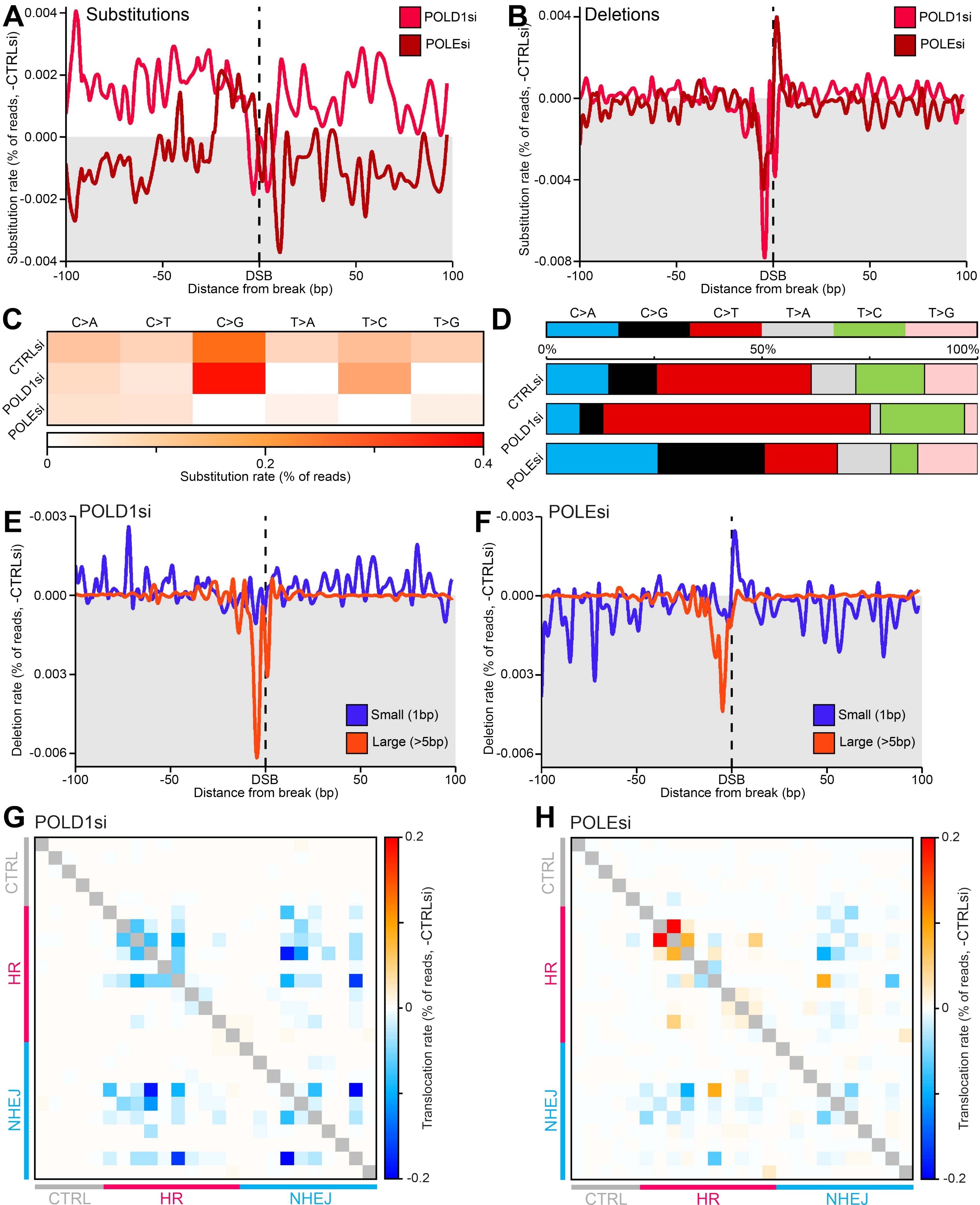

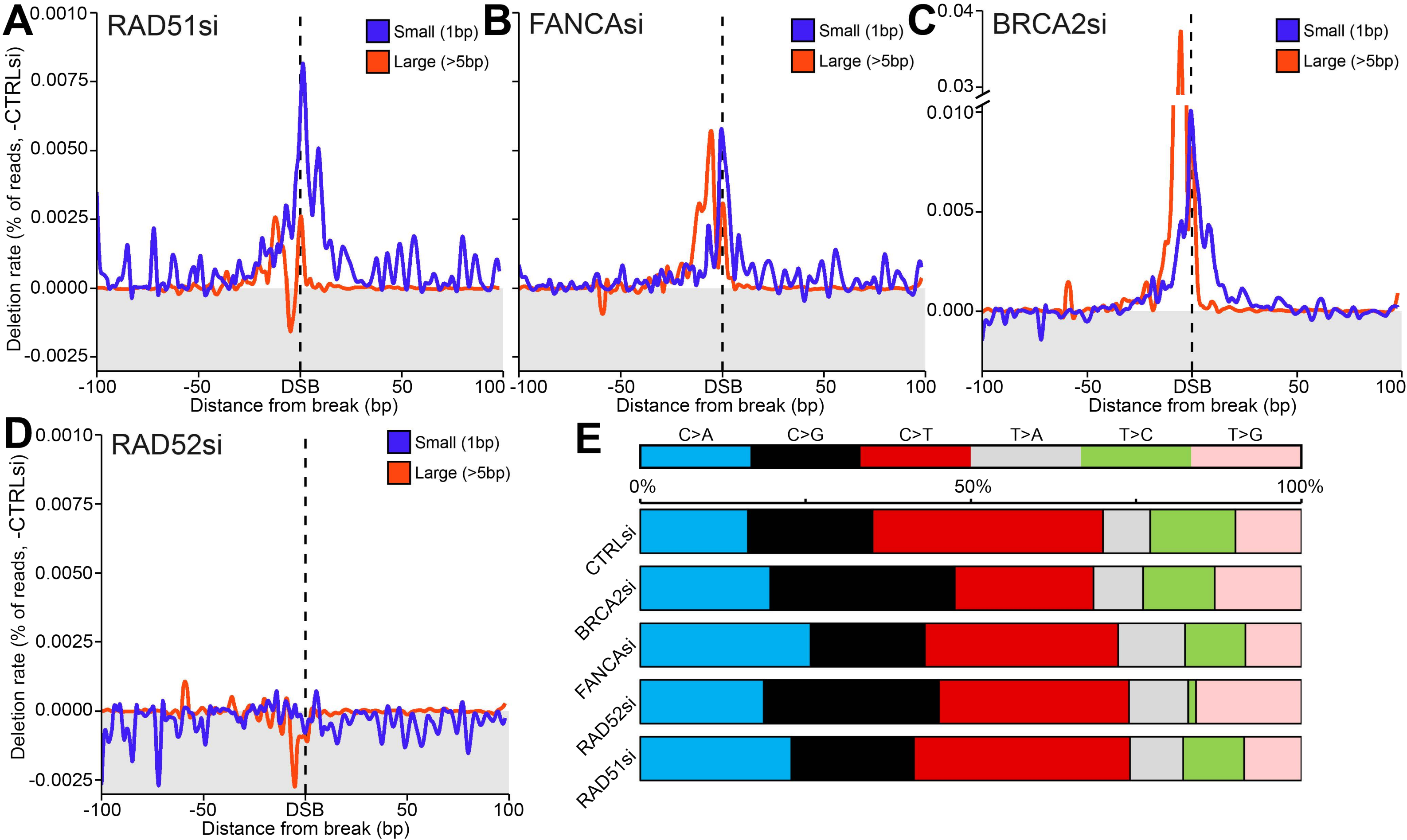
